# The inherent multidimensionality of temporal variability: How common and rare species shape stability patterns

**DOI:** 10.1101/431296

**Authors:** Jean-François Arnoldi, Michel Loreau, Bart Haegeman

## Abstract

Empirical knowledge of ecosystem stability and diversity-stability relationships is mostly based on the analysis of temporal variability of population and ecosystem properties. Variability, however, often depends on external factors that act as disturbances, making it difficult to compare its value across systems and relate it to other stability concepts. Here we show how variability, when viewed as a response to stochastic perturbations, can reveal inherent stability properties of ecological communities, with clear connections with other stability notions. This requires abandoning one-dimensional representations, in which a single variability measurement is taken as a proxy for how stable a system is, and instead consider the whole set of variability values associated to a given community, reflecting the whole set of perturbations that can generate variability. Against the vertiginous dimensionality of the perturbation set, we show that a generic variability-abundance pattern emerges from community assembly, which relates variability to the abundance of perturbed species. As a consequence, the response to stochastic immigration is governed by rare species while common species drive the response to environmental perturbations. In particular, the contrasting contributions of different species abundance classes can lead to opposite diversity-stability patterns, which can be understood from basic statistics of the abundance distribution. Our work shows that a multidimensional perspective on variability allows one to better appreciate the dynamical richness of ecological systems and the underlying meaning of their stability patterns.

## Introduction

Ecological stability is a notoriously elusive and multi-faceted concept [7, 28]. At the same time, understanding its drivers and relationship with biodiversity is a fundamental, pressing, yet enduring challenge for ecology [8, 23, 24, 26]. The temporal variability of populations or ecosystem functions, where lower variability is interpreted as higher stability, is an attractive facet of ecological stability, for several reasons. First, variability is empirically accessible using simple time-series statistics [36]. Second, variability – or its inverse, invariability – is a flexible notion that can be applied across levels of biological organization [15] and spatial scales [40, 41]. Third, variability can be indicative of the risk that an ecological system might go extinct, collapse or experience a regime shift [32]. During the last decade, the relationship between biodiversity and ecological stability has thus been extensively studied empirically using invariability as a measure of stability [5, 13, 16, 20, 27, 35].

In a literal sense, stability is the property of what tends to remain unchanged [29]. Variability denotes the tendency of a variable to change in time, so that its inverse, invariability, fits this intuitive definition. However, variability is not necessarily an inherent property of the system that is observed (e.g., a community of interacting species), as it typically also depends on external factors that act as perturbations, and generate the observed variability. In other words, the variability of a community is not a property of that community *alone*. It may be caused by a particular perturbation regime so that a different regime could lead to a different value of variability. Stronger perturbations will generate larger fluctuations, and the way a perturbation’s intensity is distributed and correlated across species is also critical. In other words, a variability measurement reflects the response of a system to the specific environmental context in which it is embedded.

Despite this complexity, quantifying the fluctuations of an ecosystem property (e.g., primary production) can be of foremost practical interest as it provides a measure of predictability in a given environmental context [11]. However, to generalize results beyond the specific context in which variability is measured, use variability to compare the stability of different systems, establish links between different stability notions, or reconcile the conflicting diversity-stability patterns and predictions reported in the empirical and theoretical literature [18], one needs to know how variability measurements can reflect a system’s inherent dynamical features.

Here, we adopt an approach in which stability is viewed as the inherent ability of a dynamical system to endure perturbations (Fig. 1A). For simplicity we will restrict to systems near equilibrium, by opposition to, e.g., limit cycles or chaotic attractors. We propose that a measure of stability should reflect, not a particular perturbation (as in Fig. 1B), but a system’s propensity to withstand *a whole class* of perturbations. We therefore consider a vast perturbation set, and study the corresponding range of community responses (Fig. 1C). Even from a theoretical perspective, considering all possible perturbations that an ecosystem can face is a daunting task. We will thus restrict our attention to model ecological communities near equilibrium, perturbed by weak stochastic perturbations, and derive analytical formulas for two complementary features of the set of their variability values: its average and maximum, corresponding to the mean-and worst-case perturbation scenarios, respectively.

**FIG. 1:**
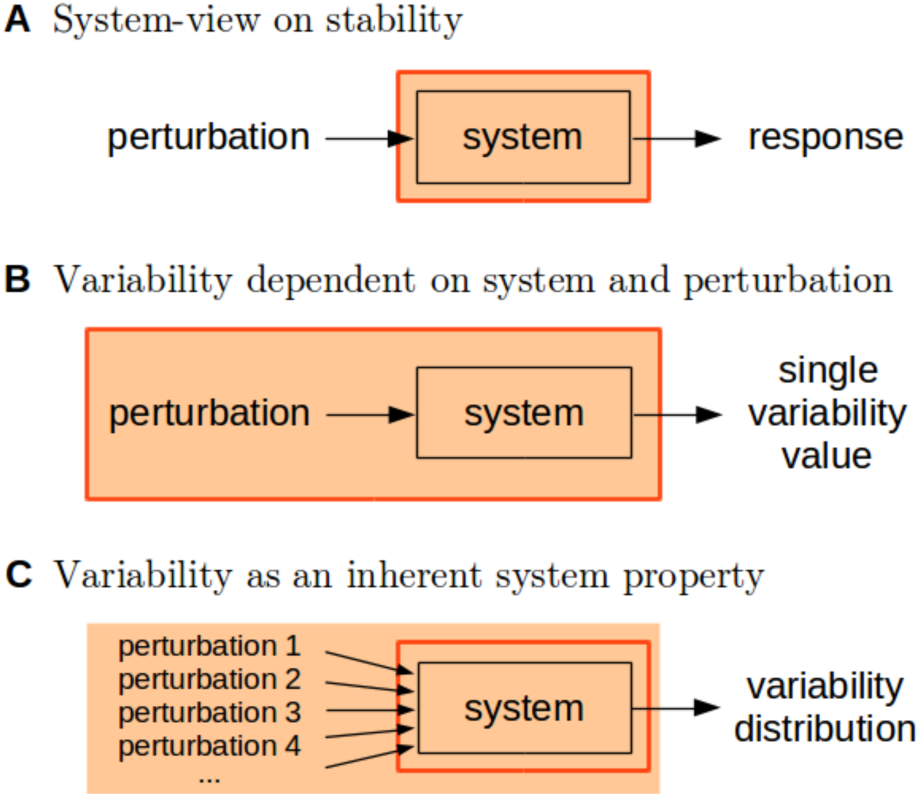
Variability vs stability. A: Stability quantifies the way a system responds to perturbations, seen as an inherent property of the system (indicated by the red framed box). B: By contrast, temporal variability is typically a feature of both the system studied and external factors that act as perturbations. C: For variability to be an inherent property of the system, one can consider a whole set of perturbations, thus integrating out the dependence on specific external factors.

After having developed a general theory of variability that can be applied to any model community near equilibrium, we turn our attention to species-rich communities that are assembled from nonlinear dynamics. We show that a generic variability-abundance pattern can emerge from the complex interactions between species during assembly. We argue that this pattern, in conjunction with the type of perturbations considered (environmental, demographic, or caused by stochastic immigration), determines the specific species abundance class that governs the variability distribution. In particular, we establish a generic link between rare species, worst-case variability, and asymptotic resilience – the long-term rate of return to equilibrium following a pulse perturbation. We finally illustrate that the contrasting contributions of various species abundance classes can be responsible for opposite diversity-invariability patterns.

The goal of our work is (i) to demonstrate that variability is an inherently multidimensional notion, reflecting the multidimensionality of an ecosystem’s responses to perturbations; (ii) to show that clear patterns exist in ecosystem responses to perturbations, which reflect the dynamical properties of distinct species abundance classes; (iii) to argue that, in order to compare and predict variability patterns, it is paramount to first identify to which abundance class these patterns or predictions refer to; and finally, (iv) to propose that a multidimensional perspective on variability allows one to better appreciate the dynamical richness of ecosystems, and the underlying meaning of their stability patterns.

### Conceptual framework

We focus on communities modelled as dynamical systems at equilibrium, and study their responses to a whole class of stochastic white-noise forcing. In this section we outline the theory, focusing on ecological intuitions, while Appendix A through D provides a self-contained presentation of its mathematical foundations. Our work follows traditional approaches of theoretical ecology [19, 24], extending the analysis to encompass a large perturbation set.

### Perturbed communities

Let *N*_*i*_ (*t*) represent the abundance (or biomass) of species *i* at time *t*, and *x*_*i*_ (*t*) = *N*_*i*_ (*t*) − *N_i_* its displacement from an equilibrium value *N*_*i*_, with *i* running over *S* coexisting species that form an ecological community. We model fluctuations in abundance (hence variability) as a response to some stochastic forcing. We focus on stationary fluctuations caused by weak perturbations with zero mean (cf. note [43]), which are governed by the following dynamical system, written from the perspective of species *i* as

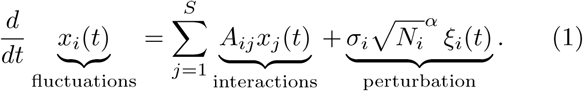

In this expression, the coefficients *A*_*ij*_ represent the effect that a small change of abundance of species *j* has on the abundance of species *i*. Organized in the *community matrix A* = (*A*_*ij*_), they encode the linearization of the nonlinear system of which (*N*_*i*_) is an equilibrium. In the perturbation term, *ξ*_*i*_ (*t*) denotes a standard white-noise source [1, 38]. In discrete time *ξ*_*i*_ (*t*) would be a normally distributed random variable with zero mean and unit variance, drawn independently at each time step (Appendix A).

Community models of the form eq. (1) were studied by Ives et al. [19] to analyze ecological time series. In their approach, stability properties are inferred from the system’s response to specific perturbations. Here we build on a similar formalism, but explicitly explore a vast set of possible perturbations. Although environmental fluctuations often follow temporal patterns [10, 31, 39] we will not consider autocorrelated perturbations. It would thus be interesting to extend the analysis to more general temporal structures of perturbations, as well as to nonlinear behaviors. What we will explicitly consider, however, are temporal correlations between *ξ*_*i*_ (*t*) and *ξ*_*j*_ (*t*), a situation in which individuals of species *i* and *j* are similar in their perception of a given perturbation, a property known to have potentially strong, and unintuitive effects on species dynamics [30].

For the fluctuations of species abundance in eq. (1) to be stationary, the equilibrium state (*N*_*i*_) must be stable. More technically, the eigenvalues of the community matrix *A* must have negative real part [14, 24]. The maximal real part determines the slowest long-term rate of return to equilibrium following a pulse perturbation. This rate is a commonly used stability measure in theoretical studies; we call it *asymptotic resilience* and denote it by ℛ_∞_ [4]. To illustrate the connections between stability concepts, we will compare asymptotic resilience to measures of variability.

### Perturbation type

The perturbation term in eq. (1) represents the direct effect that a perturbation has on the abundance of species *i*. It consists of two terms: some power *α* of 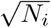, and a species-specific term *σ*_*i*_ *ξ*_*i*_ (*t*). The latter is a function of the perturbation itself, and of traits of species *i* that determine how individuals of that species perceive the perturbation. The former defines a statistical relationship between a perturbation’s direct effects and the mean abundance of perturbed species. It allows us to consider ecologically distinct sources of variability (Fig. 2).

**FIG. 2:**
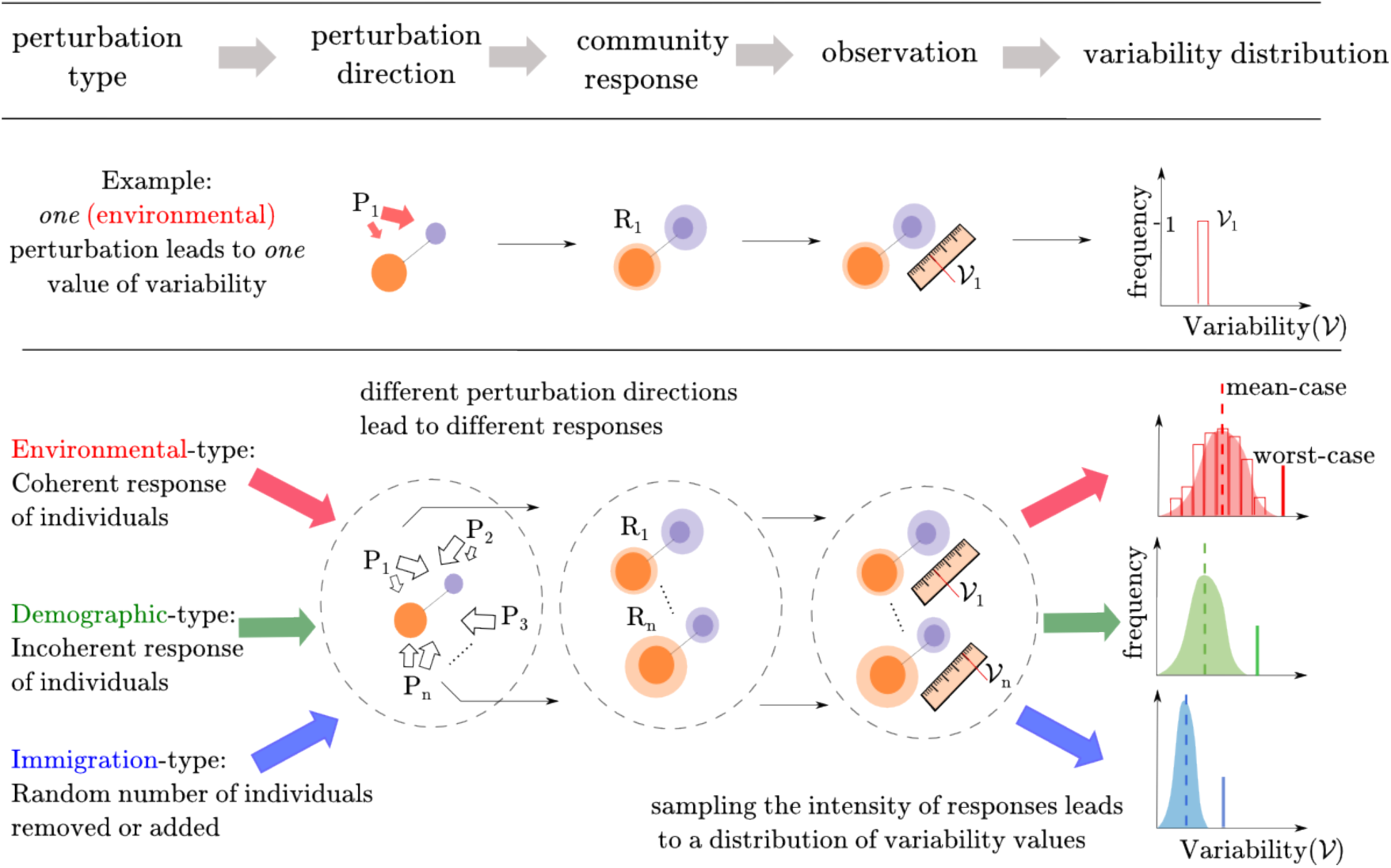
A theoretical framework for variability. Perturbations are characterized by their type, a statistical relationship between the direct effect of perturbations and the abundance of perturbed species. For a given type and fixed intensity, there remains a whole set of covariance structures of perturbations, i.e., various perturbation directions, that will be transformed by community dynamics into a whole set of community responses, i.e., various covariance structures of species stationary time series. A sampling of those responses leads to a variability distribution, one for each perturbation type. Spanning all perturbation types leads to a family of variability distributions (in blue, green and red in the rightmost column).

When individuals of a given species respond in synchrony to a perturbation, the direct effect of the perturbation will be proportional to the abundance of the perturbed species, thus a value of *α* close to 2 [22]. We call this type of perturbation *environmental* as fluctuations of environmental variables typically affect all individuals of a given species, leading, e.g. to changes in the population growth rate [25].

If individuals respond incoherently, e.g., some negatively and some positively, the direct effect of the perturbation will scale sublinearly with species abundance. For instance, demographic stochasticity can be seen as a perturbation resulting from the inherent stochasticity of birth and death events, which are typically assumed independent between individuals. In this case *α* = 1, and we thus call such type *demographic* [22].

We can also consider purely exogenous perturbations, such as the random removal or addition of individuals. In this case *α* = 0. We call such perturbations *immigration*-type but stress that actual immigration events do not necessarily statisfy this condition (e.g., they can be density-dependent). Furthermore, because we focus on zero-mean perturbations, perturbations of this type contain as much *emigration* than immigration. The reasoning behind this nomenclature is that, in an open system, fluctuations of an otherwise constant influx of individuals (immigration flux) would correspond to an immigration-type perturbation.

More generally, eq. (1) with *α* ∈ [0, 2] can describe a continuum of perturbation types. Note that, although not unrelated, such a statistical relationship between a perturbation’s direct effects and the abundance of perturbed species is not equivalent to Taylor’s law [33]. The latter is an empirically observed power-law relationship between the variance and mean of population fluctuations. Hence, in contrast to the perturbation type *α*, the exponent of Taylor’s law depends on community dynamics, e.g., on species interactions [21]. We will come back to this point below and in the Discussion.

### Perturbation intensity

For a given community, a stronger perturbation will lead to stronger fluctuations. A disproportionate increase in their amplitude as perturbation intensity changes would reveal nonlinearity in the dynamics [42]. In a linear setting, however, such effects cannot occur and there is only a linear dependency on perturbation intensity. This trivial dependency can be removed by controlling for perturbation intensity. We now illustrate how to do so, for a given definition of variability.

In our setting, fluctuations induced by white-noise forcing are normally distributed, thus fully characterized by their variance and covariance. It is therefore natural to construct a measure of variability based on the variance of species time-series. To compare variability of communities with different species richness we will measure their average variance:

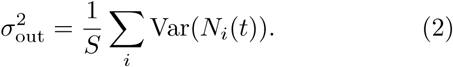

In empirical studies, variability is often associated to an ecosystem function (primary productivity, ecosystem respiration, etc). This amounts to measuring the ecosystem response along a direction in the space of dynamical variables. In Appendix B we explain how considering the average variance amounts to taking the expected variance along a random choice of direction of observation. In this sense, eq. (2) represents the variance of a “typical” observation.

We now wish to remove the trivial effect of perturbation intensity from eq. (2). Let us start from a one-dimensional system *dx/dt* = −*λ x* + *σ ξ* (*t*). Its stationary variance is 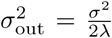. Here we see the combined effect of perturbation (*σ* ^2^ and dynamics *λ*, leading us to define (*σ* ^2^ as measure of perturbation intensity. For species-rich communities, we define perturbation intensity as the average intensity per species, that is, using the species-specific intensities 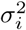:

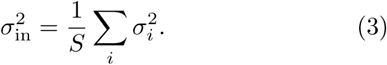

When increasing all species-specific perturbation intensities by a factor *c*, both 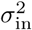 and 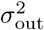 increase by the same factor. To remove this linear dependence, we define vari-ability as

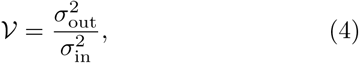

i.e., as the average species variance relative to perturbation intensity (see 19 for a similar definition of variability). Generalizing previous work [3, 4] to an arbitrary perturbation type, we construct *invariability* as

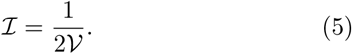

The factor 1/2 allows ℐ to coincide, for simple systems, with asymptotic resilience [4]. In particular, for the one-dimensional example considered above for which ℛ_∞_ =λ, we do have 𝒱 = 1/2λ and thus ℐ = λ= ℛ_∞_.

### Perturbation direction

Once intensity 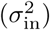 is controlled for, perturbations can still differ in how their intensity is distributed 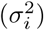 and correlated in time (correlation between *ξ*_*i*_ (*t*) and *ξ*_*j*_ (*t*), see eq. (1)) across species. We want to be able to model the fact that species with similar physiological traits will be affected in similar ways by, say, temperature fluctuations, whereas individuals from dissimilar species may react in unrelated, or even opposite, ways [30]. We will thus study the effect of the covariance structure of the perturbation terms, i.e., the effect of the *direction* of perturbations. Spanning the set of all perturbation directions will define a whole range of community responses. Assuming some probability distribution over this set translates into a probability distribution over the set of responses, i.e., a variability distribution (see Fig. 2). We will assume all perturbation directions to be equiprobable, but our framework allows different choices of prior. Finally, spanning the set of perturbation types reveals a continuous *family* of variability distributions. In Fig. 2 we show three archetypal elements of this family, corresponding to *α* = 0 (blue distribution), *α* = 1 (green distribution) and *α* = 2 (red distribution).

For each distribution we consider two complementary statistics: mean-and worst-case responses. In Appendices C and D we prove that the worst-case response is always achieved by a perfectly coherent perturbation, i.e., a perturbation whose direct effects on species are not independent, but on the contrary, perfectly correlated in time. We derive explicit formulas to compute the worst-case variability from the community matrix and species equilibrium abundances, see eqs. (C2, D5). The mean-case scenario, on the other hand, is defined with respect to a prior over the set of perturbation directions. For the least informative prior, we prove in Appendices C and D that a perturbation affecting all species independently but with equal intensity, realizes the mean-case response. This provides a way to compute this response from the community matrix and the species abundances, given in eqs. (C3, D6).

### Variability patterns for two-species community

Before considering complex communities, let us illustrate our variability framework on the following elementary example, in the form of a 2 × 2 community matrix

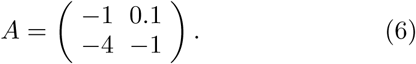

This matrix defines a linear dynamical system that could represent a predator-prey community, with the first species benefiting from the second at the latter’s expense. Its asymptotic resilience is ℛ_∞_ = 1. Let us suppose that the prey, *N*_2_ (second row/column of *A*) is 7.5 times more abundant than its predator, *N*_1_ (first row/column of *A*) and consider stochastic perturbations of this community, as formalized in eq. (1).

In Fig. 3 we represent the set of perturbation directions as a disc, in which every point is a unique perturbation direction (see Appendix E for details). The effect of a perturbation on a community is represented as a color; darker tones imply larger responses, with the baseline color (blue, green or red) recalling the perturbation type (α = 0, 1, 2, respectively). Points at the boundary of the disc correspond to coherent perturbations, which have the potential to generate the largest (but also the smallest) variability. This is why the color maps of Fig. 3 take their extreme values at the boundary. We see that variability strongly depends on the perturbation direction, and that this dependence is strongly affected by the perturbation type. For immigration-type perturbations (in blue) variability is largest when perturbing the predator species most strongly (the least abundant species in this example). For demographic-type perturbations (in green) perturbations that equally affect the two species but in opposite ways achieve the largest variability. For environmental-type perturbations (in red) variability is largest when perturbing the prey species (the most abundant species in this example). For all types we see that positive correlations between the components of the perturbation (i.e., moving upwards on the disc) reduce variability (see 30 for related results).

**FIG. 3:**
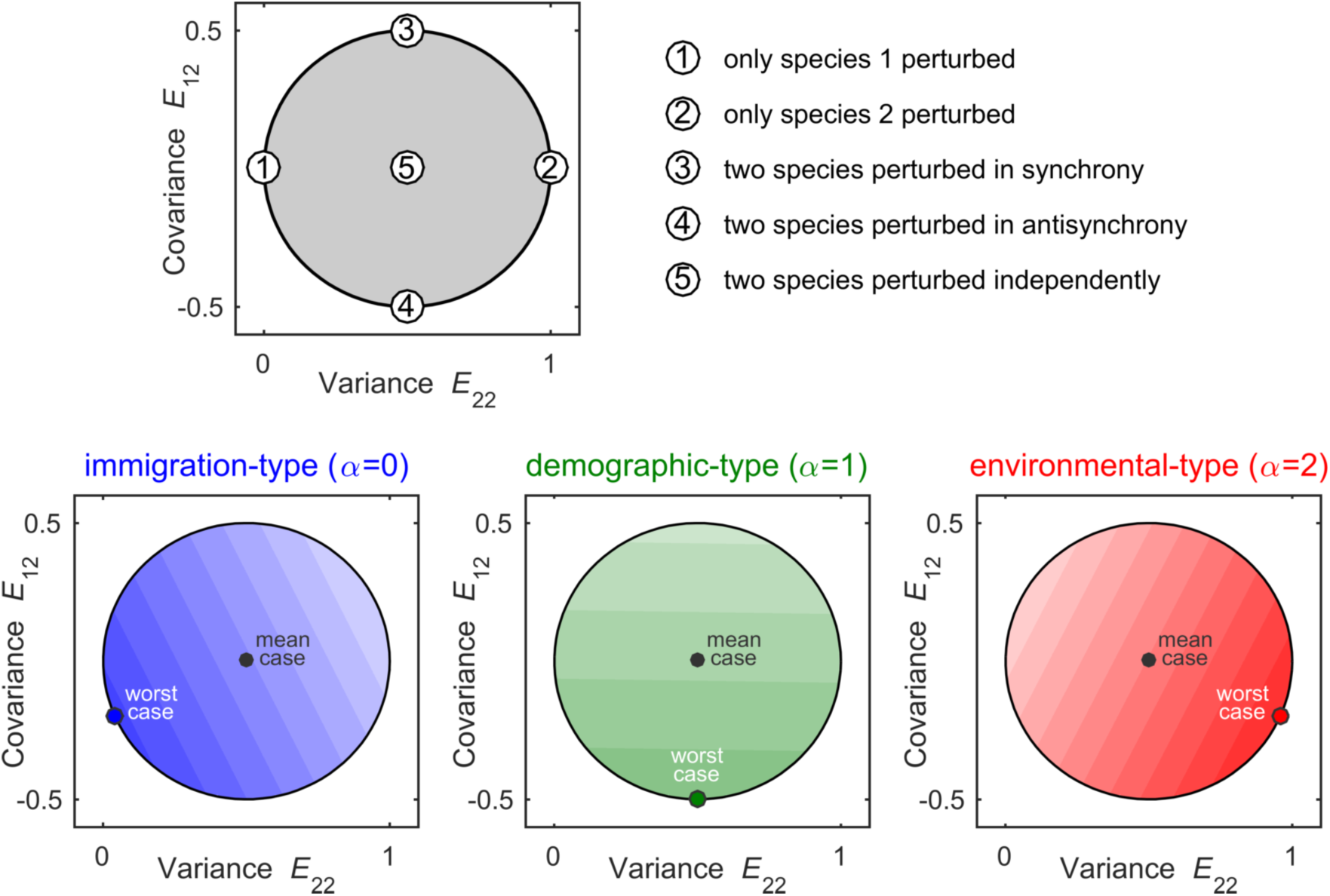
Variability patterns for a two-species community. Top panel: For a two-species community the set of all perturbation directions can be represented graphically as a disc (shaded in gray), with the variance of the perturbation term *ξ*_*2*_ (*t*) on the *x*-axis and the covariance between *ξ*_*1*_ (*t*) and *ξ*_*2*_ (*t*) on the y-axis. Some special perturbation directions are indicated (numbers 1 to 5, see also Appendix E). Bottom panels: We consider a predator-prey system; the community matrix *A* is given by eq. (6), and the prey (species 2) is 7.5 more abundant than its predator (species 1). The induced variability depends on the perturbation directions (darker colors indicate larger variability), and this dependence in turn depends on the perturbation type *α.* For immigration-type perturbations *(α* = 0, in blue) variability is largest when perturbing species 1 most strongly. For demographic-type perturbations (*α* = 1, in green) perturbations that affect the two species equally strongly but in opposite ways achieve the largest variability. For environmental-type perturbations (*α* = 2, in red) variability is largest when perturbing species 2 most strongly. Notice that the worst case is always achieved by perturbations lying on the edge of the perturbation set. Such perturbations are perfectly correlated (see main text and Appendix E).

Thus, in general, a given community cannot be associated to a single value of variability. Depending on the type of perturbations causing variability, different species can have completely different contributions. This stands in sharp contrast with asymptotic resilience ℛ_∞_, which associates a single stability value to the community. Although we know from previous work [4] that the smallest invariability value in response to immigration-type perturbations will always be smaller than ℛ_∞_, in general (i.e., any perturbation type and/or any perturbation direction) there is, a priori, no reason to expect a relationship between invariability and asymptotic resilience.

### Variability patterns in complex communities

The dimensionality of variability will be larger in communities comprised of many species, as their sheer number, *S*, increases the dimension of the perturbation set quadratically. Yet, when species interact, a generic structure can emerge from ecological assembly, revealing a simple relationship between variability and the abundance of perturbed species. To show this, we study randomly assembled communities. We start from a large pool of species with randomly drawn dynamical traits (cf. note [44]), and let the system settle to an equilibrium following Lotka-Volterra dynamics. During assembly some species would go extinct, but no limit cycles, chaotic behavior or multi-stability were observed. A complete description of the nonlinear model is given in Appendix F and Matlab simulation code is available as supplementary material.

In Fig. 4 we show the variability patterns for a single randomly assembled community, but the results hold more generally (see below). The species pool consisted of *S*_pool_ = 50 species, with species interaction strengths an order of magnitude smaller than the strengths of species self-regulation (see Appendix F for details). The assembled community had *S* = 40 coexisting species. In this species-rich context, the perturbation set cannot be represented exhaustively. We therefore plot the variability induced by species-specific perturbations (of various types) against the abundance of perturbed species. That is, we focus on the effect of a specific subset of perturbations, those affecting a single species. Linear combinations of these perturbations will span all scenarios in which species are affected independently, but exclude scenarios in which they are perturbed in systematically correlated or anti-correlated way (cf. note [45]).

**FIG. 4:**
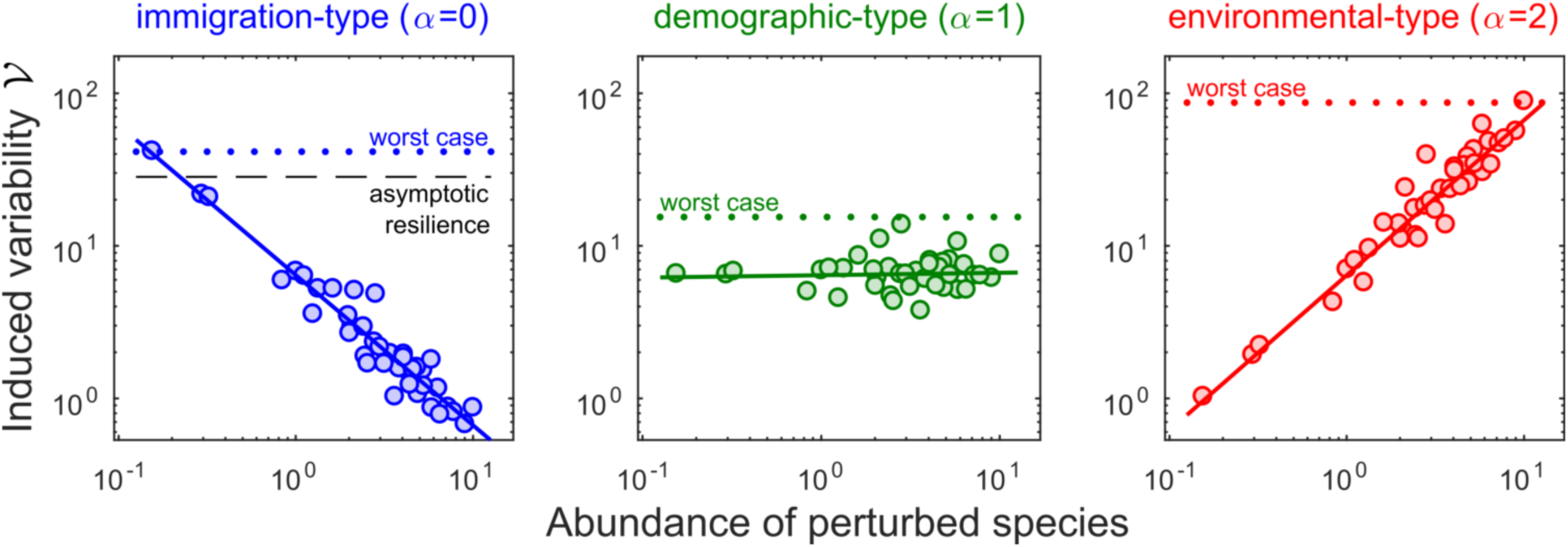
Variability-abundance pattern in a complex community. We consider a community of *S* = 40 species, and look at the variability induced by perturbing a single species, whose abundance is reported on the x-axis. Left: When caused by immigration-type perturbations (*α* = 0), variability is inversely proportional to the abundance of the perturbed species (notice the log scales on both axis). The worst case is achieved by perturbing the rarest species, and is determined by asymptotic resilience (more precisely, it is close to 1/2 ℛ_∞_). Middle: For demographic-type perturbations (*α* = 1), variability is independent of the abundance of the perturbed species. The worst case is not necessarily achieved by focusing the perturbation on one particular species. Right: For environmental-type perturbations (*α* = 2), variability is directly proportional to the abundance of the perturbed species. The worst case is attained by perturbing the most abundant.

The leftmost panel shows a negative unit slope on log scales: when caused by immigration-type perturbations, variability is inversely proportional to the abundance of perturbed species. Randomly adding and removing individuals from common species generates less variability than when the species is rare. In fact, the worst-case scenario corresponds to perturbing the rarest species. Worst-case invariability is close to asymptotic resilience, which corroborates previous findings showing that the long-term rate of return to equilibrium is often associated to rare species [2, 15]. On the other hand, the middle panel of Fig. 4 shows that, in response to demographic-type perturbations, variability is independent of perturbed species’ abundance. Finally, the right-most panel shows a positive unit slope on log scales: when caused by environmental-type perturbations, variability is proportional to the abundance of perturbed species. The worst case is thus attained by perturbing the most abundant one. Despite being more stable than rare ones (they buffer exogenous perturbations more efficiently, see left-hand panel), common species are more strongly affected by environmental perturbations, and can thus generate the most variability.

Those patterns are not coincidental, but emerge from species interactions, as we illustrate in Fig. 5. In their absence, other patterns can be envisioned. This is because, without interactions, the response to a species-specific perturbation involves the perturbed species only. The variability-abundance relationship is then 𝒱 = *N*^*α*^ /2*r*, with *N* = *K*. If *r* and *K* are statistically independent in the community (top-left panel in Fig. 5), this yields a different scaling than the one seen in Fig. 4. In the case of an r-K trade-off (i.e., species with larger carrying capacities have slower growth rate), abundant species would be the least stable species (bottom-left panel in Fig. 5, in blue) which is the opposite of what the leftmost panel of Fig. 4 shows. However, as interaction strength increases (from left to right in Fig. 5; the ratios of inter-to intraspecific interaction strength are 0, 0.02 and 0.1 approximately), we see emerging the relationship between abundance and variability of Fig. 4, regardless of the choice made for species growth rates and carrying capacities. We explain in Appendix G why this reflects a generic, limit-case behavior of complex communities. It occurs when species abundances, due to substantial indirect effects during assembly, become only faintly determined by their carrying capacities (cf. note [46]). Importantly, our example demonstrates that this limit can be reached even for relatively weak interactions (in Fig. 4 and in the right-hand panels of Fig. 5, the interspecific interaction strengths are ten times smaller than the intraspecific ones).

**FIG. 5:**
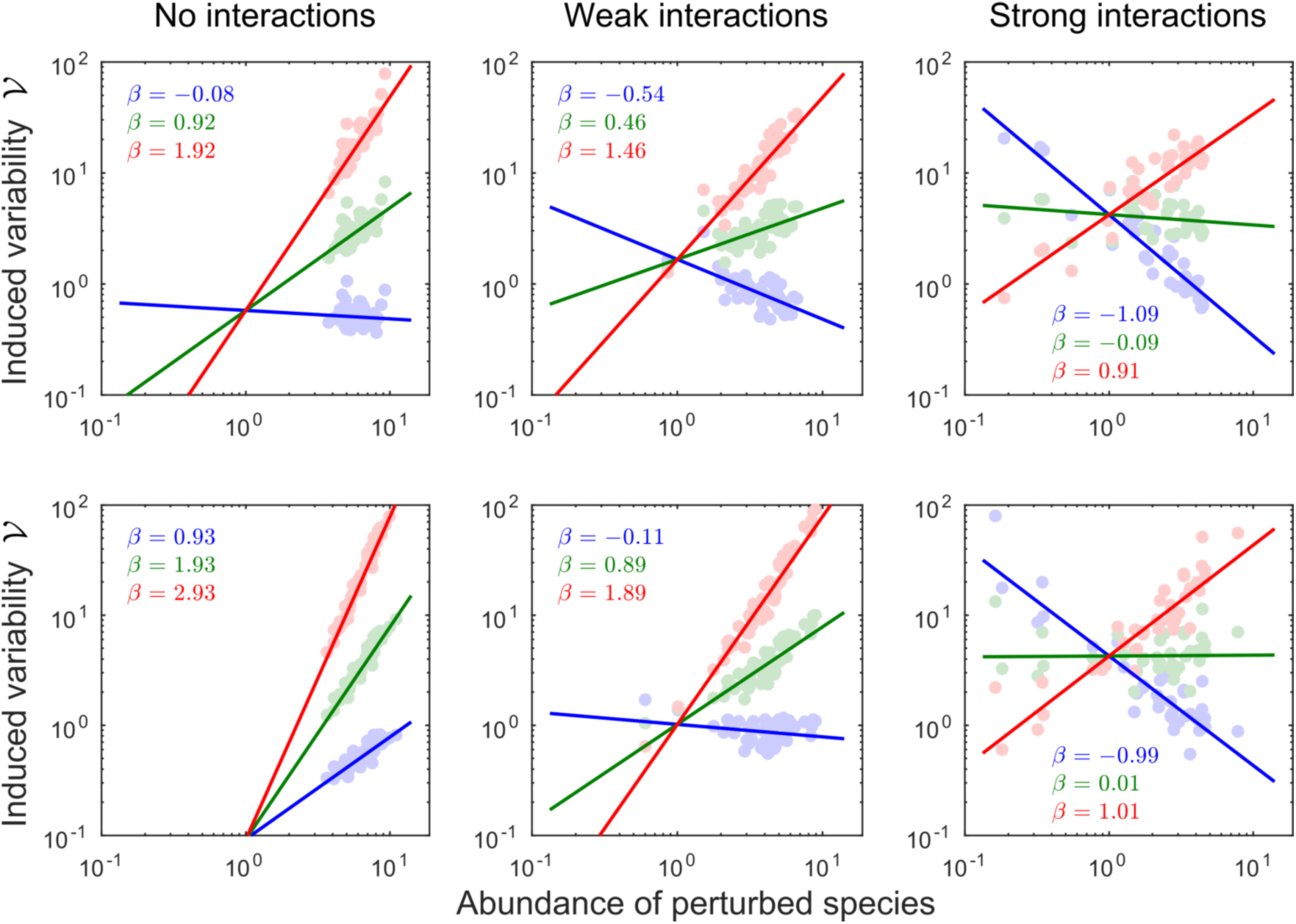
The emergence of the variability-abundance pattern (same procedure as in Fig. 4). Top row: intrinsic growth rates *r* and carrying capacities *K* are sampled independently. Bottom row: Species satisfy a *r*-*K* trade-off (*r* ∼ 1/*K*). Colors correspond to the three perturbation types: *α* = 0 (blue), *α* = 1 (green) and *α* = 2 (red). The value /3 reported in each panel corresponds to the exponent of the fitted relationship 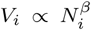 for each perturbation type. As interaction strength increases (left to right) we see emerging the relationship between abundance and variability described in Fig. 4, i.e., *β* = *α* - 1. Thus when species interactions are sufficiently strong, variability always ends up being: (blue) inversely proportional, (green) independent and (red) directly proportional to the abundance of the perturbed species. Note that such relationships differ from Taylor’s law: they represent an average community response to individual species perturbations, whereas Taylor’s law deals with individual species responses to a perturbation of the whole community.

Although we considered a specific section of the perturbation set, the response to single-species perturbations of immigration and environmental types can still span the whole variability distribution, from worst-case (rarest and most abundant species perturbed, respectively) to mean-and best-case scenarios (most abundant and rarest species perturbed, respectively). For demographic-type perturbation the situation is more subtle as the response is independent of species abundance, and, in general, extreme scenarios will be associated to temporally corre-lated perturbations affecting multiple species.

The variability-abundance patterns shown in Figs. 4 and 5 should not be confused with Taylor’s law [33], a power-law relationship between a species’ variance and its mean abundance. In fact, the variability-abundance pattern is *dual* to Taylor’s law, it represents the community response to single-species perturbations instead of that of individual species to a community-wide perturbation (cf. note [47]).

### Diversity-invariability relationships

To illustrate some implications of the generic variability-abundance pattern, we now propose to revisit the diversity-stability relationship, with stability quantified as invariability ℐ. For a given size of the species pool, we randomly sample species dynamical traits to assemble a stable community. By increasing the size of the pool we generate communities of increasing species richness *S*. For each community, we uniformly sample the boundaries of its perturbation set by drawing 1000 fully correlated perturbations (i.e., those that can realize the maximal response), of a given type. We compute the bulk of the resulting invariability distribution (5 to 95 percentiles), as well as its mean and extreme realized values. We also compute theoretical predictions for mean-and worst-case scenarios, and asymptotic resilience ℛ_∞._

The leftmost panel of Fig. 6 shows a negative relation-ship between immigration-type invariability and species richness. Asymptotic resilience and worst-case invariabil-ity mostly coincide, with a decreasing rate roughly twice as large as that of the mean case. The middle panel suggests a different story. Mean-case demographic-type invariability stays more or less constant whereas the worst case diminishes with species richness, although much more slowly than ℛ_∞_. The relationship between diversity and stability is thus ambiguous. In the rightmost panel we see an increase in all realized environmental-type invariability values with species richness, showcasing a positive diversity-stability relationship.

**FIG. 6:**
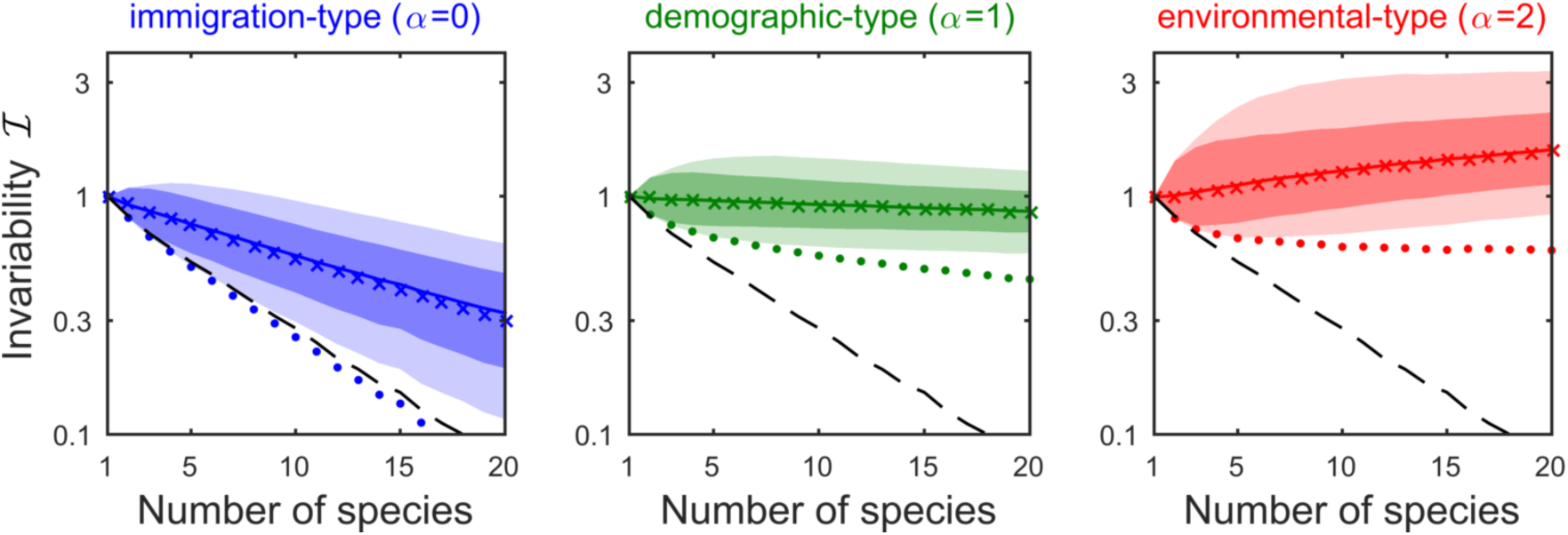
Different perturbation types yield contrasting diversity-stability relationships, with stability quantified as invariability ℐ. We generated random communities of increasing species richness *S* and computed their invariability distribution in response to 1000 random perturbations. Full line: median invariability, dark-shaded region: 5th to 95th percentile, light-shaded region: minimum to maximum realized values. The × -marks correspond to the analytical approximation for the median, the dots to the analytical formula for the worst-case. Dashed line is asymptotic resilience ℛ_∞_. For immigration-type perturbations (*α* = 0, blue) diversity begets instability, with ℛ_∞_ following worst-case invariability. For demographic-type perturbations (*α* = 1, green) the trend is ambiguous. For environmental-type perturbations (*α* = 2, red) all realized values of invariability increase with *S*.

The different relationships between diversity and stability can be understood in terms of the generic variability-abundance patterns of Figs. 4 and 5 (see Ap-pendix H for details). In the case of immigration-type variability, species contributions to variability are proportional to the inverse of their abundance (first panel of Fig. 4). Hence, the worst-case scenario follows the abundance of the rarest species, which rapidly declines with species richness. As detailed in Appendix H, mean-case invariability scales as the average species abundance, which also typically decreases with *S*.

The responses to demographic perturbations, on the other hand, are not determined by any specific species abundance class (second panel of Fig. 4), so that no simple expectations based on typical trends of abundance distributions can be deduced.

We recover a simpler behavior when looking at the response to environmental-type perturbation. It is now abundant species that drive variability (rightmost panel of Fig. 4). As explained in Appendix H, mean-case invariability now scales as the *inverse* of an average species abundance. The latter typically declines with *S* explaining the observed increase of mean-case invariability.

In all panels of Fig. 6, the bulk of invariability stays close to the meanwhile moving away from the worst-case. This is because the worst-case corresponds to a single direction of perturbation met with the strongest response, a fine-tuned perturbation which becomes increasingly unlikely to be picked at random as S increases. There is an analogy to be made between stability and diversity. As has been said about diversity metrics (e.g., species richness, Simpson index or Shannon entropy), different invariability measures *“differ in their propensity to include or to exclude the relatively rarer species”* [17]. In this sense, different invariability measures can probe different dynamical aspects of a same community, with potentially opposite dependencies on a given ecological parameter of interest.

## Discussion

Because it is empirically accessible using simple time-series statistics, temporal variability is an attractive facet of ecological stability. But there are many ways to define variability in models and empirical data, a proliferation of definitions reminiscent of the proliferation of definitions of stability itself [12]. Variability measurements often depend, not only on the system of interest, but also on external factors that act as disturbances, which makes it difficult to relate variability to other stability concepts. These caveats constitute important obstacles toward a synthetic understanding of ecological stability, and its potential drivers [18].

We proposed to consider variability as a way to probe and measure an ecosystem’s response to perturbations, thus revealing inherent dynamical properties of the perturbed system. We did not seek for an optimal, single measure of variability but, on the contrary, we accounted for a vast set of perturbations, leading to a whole distribution of responses. We focused on the worst-and mean-case values of this distribution as functions of species abundance, their interactions, and the *type* of perturbations that generates variability.

A perturbation type characterizes a statistical relationship between its direct effect on a population and the latter’s abundance. We distinguished between environmental perturbations, whose direct effects on populations scales proportionally to their abundance; demographic perturbations, whose direct effect on populations scales sublinearly to their abundance; and purely exogenous perturbations, representing random addition and removal of individual, independent of the size of the perturbed population (immigration-type). Controlling for perturbation type and intensity, we considered all the ways this intensity can be distributed and correlated across species.

After having described a general (linear) theory for variability, which emphasizes its highly multidimensional nature, we turned our attention towards species-rich communities assembled by random (nonlinear) Lotka-Volterra dynamics. Because of the sheer number of species contained in such communities (*S* ≈40 in our examples), we could have expected the dimensionality of perturbations and responses to be so large that variability distributions would be too complex and broad to be clearly described. However, the process of assembly allowed for a simple behavior to emerge: a generic relationship between variability and the abundance of individually perturbed species. In essence, this pattern predicts that the species’ ability to buffer exogenous perturbations is proportional to their abundance. In conjunction to this simple pattern, the type of perturbation will then determine the individual contributions of species to the variability distribution, so that both common and rare species can determine variability. This is reminiscent of diversity measures [17], some of which (e.g., species richness) are sensitive to the presence of rare species, while others are mostly indicative of the distribution of abundant species (e.g., Simpson diversity index).

These connections with different diversity metrics can explain contrasting trends in invariability as a function of species richness. Since immigration-type perturbations mostly affect rare species, they lead to a negative diversity-invariability relationship, reflecting a growing number and rarity of rare species. On the other hand, in response to demographic perturbations, species contributions to variability can be independent of their abundance. In this case, variability is not expected to follow trends in diversity, so that diversity-invariability patterns can be less predictable and harder to interpret. Finally, although common species buffer exogenous perturbations efficiently, they are also the most affected by environmental-type perturbations. This can lead to a proportional relationship between average abundance and mean-case invariability, and hence to a positive diversity-invariability relationship.

### Implications for empirical patterns

Our theoretical models show wide variability distributions. These distributions would become even wider when accounting for nonlinear system dynamics and temporally autocorrelated perturbations. Therefore, we also expect a large dispersion of empirical variability data, i.e., when the variability of the same system is measured repeatedly. For certain applications it might be sufficient to restrict to a particular perturbation regime, but in order to detect in variability an inherent stability property of a system, i.e. a property that is not bound to a specific environmental context (see Fig. 1), one must describe of the spread of variability.

To do so, the most direct approach consists in observing the same community under multiple environmental conditions. With relatively few observations, one can estimate the mean and spread of the response distribution. There is, however, more information to be extracted from a time series than a single variability value. If high-quality time series are available, it might be possible to infer linear model dynamics, which can then be used to compute stability properties [19], and in particular, variability distributions.

We showed that species abundances greatly affect variability distributions. This new insight has broad consequences. For example, it has been reported that ecosystem-level and population-level stability tend to increase and decrease, respectively, with increasing diversity [5, 20]. Ecosystem-level stability is often quantified based on the variability of total biomass, which gives, by construction, a predominant weight to abundant species. On the other hand, averages of single-species variabilities have been used to measure population-level stability [34]. These averages are strongly affected, and can even be fully determined, by rare, highly variable species [15]. Thus, here as well as in our theoretical results (Fig. 6), stability metrics governed by common, or rare, species tend to generate respectively positive and negative diversity-stability relationships. It would be interesting to test whether this observation holds more generally, e.g., if it can explain the contrasting relationships recently reported by Pennekamp et al. [27].

The type of perturbations affects which species abundance class contributes most to variability. In turn, the physical size of the system considered affects which perturbation type dominates. This is well known in population dynamics [9], but it also transposes to the community level. At small spatial scales, implying small populations, we may expect variability to be driven by demographic stochasticity. At larger scales, implying larger populations, demographic stochasticity will be negligible compared with environmental perturbations. Just as changing the perturbation type transforms the respective roles of common and rare species, patterns of variability at different scales should reflect different aspects of a community [6], associated to different species abundance classes (abundant species at large spatial scales, rare/rarer species at small spatial scales).

Empirically determining the perturbation type, which is a preliminary step to test the stability patterns predicted in this paper, is a non-trivial task. To develop suitable methods, it might be helpful to first understand the link between the variability-abundance patterns (see Figs. 4 and 5) and Taylor’s law [33]. The latter is an empirically accessible pattern, relating the mean and variance of population sizes. A close connection is indeed expected: we studied the behavior of the community response to an individual species perturbation, while Taylor’s law focuses on the individual species response to a perturbation of the whole community. This duality also suggests that Taylor’s law is, at the community level, strongly affected by species interactions. Although this is known [21], our approach could shed new light on the information regarding species interactions and other dynamical traits, actually contained in community-level Taylor’s laws.

### Link with other stability measures

We noted a connection between variability and asymptotic resilience, which is a popular notion in theoretical studies [7]. We showed that asymptotic resilience is comparable to the largest variability in response to an immigration-type perturbation, which is often a perturbation of the rarest species (first panel of Fig. 4). While asymptotic resilience is sometimes considered as a measure representative of collective recovery dynamics, we previously explained why that this is seldom the case [2]. The asymptotic rate of return to equilibrium generally reflects properties of rare “satellite” species, pushed at the edge of local extinction by abundant “core” species. On the other hand, short-time return rates can exhibit qualitatively different properties related to more abundant species.

In fact, the multiple dimensions of variability are related to the multiple dimensions of return times. Variability is an integral measure of the transient regime following pulse perturbations, i.e., a superposition of responses to various pulses, some of which have just occurred and are thus hardly absorbed, while others occurred long ago and are largely resorbed. If abundant species are faster than rare ones (the case in complex communities, see Appendix G), if they are also more strongly perturbed (e.g., by environmental perturbations), the bulk of the transient regime will be short: variability in response to environmental perturbations is associated with a short-term recovery. By contrast, if all species are, on average, equally displaced by perturbations (e.g., by immigration-type perturbations), rare species initially contribute to the overall community displacement as much as do abundant ones. Since their recovery is typically very slow, the transient regime will be long: variability in response to immigration-type perturbations is associated with a long-term recovery.

Ecologists have long acknowledged the multi-faceted nature of ecological stability [7, 12, 18, 28], but here we show that a single facet (variability) is in itself inherently multidimensional, thus suggesting that links across facets can be subtle. Short-term return rates may be linked with environmental variability, but environmental variability may have nothing to do with immigration-type variability, the latter possibly related with long-term return rates and driven by rare species. Because measures can be determined by different species abundance classes, we should not expect a general and simple connection to hold between facets of ecological stability.

## Conclusion

The multidimensional nature of variability can lead to conflicting predictions, but once this multidimensionality is acknowledged, it can be used to extensively probe the dynamical properties of a community. In particular, in species-rich systems, we revealed a generic pattern emerging from ecological assembly, relating species abundance to their variability contribution. This allowed connections to be drawn between variability and statistics of abundance distributions. We argued that similar patterns should underlie ecosystem responses to other families of perturbations (e.g., pulse perturbations). Therefore, we conclude that embracing the whole set of a ecosystem responses can help provide a unifying view on ecological stability and shed new light on the meaning of empirical and theoretical stability patterns.

## Supporting information

Matlab code used to generate figures

## Supplementary material

Matlab simulation code is available online (bioRxiv server) DOI: 10.1101/431296 (supplementary material).

## Acknowledgements

We thank Matthieu Barbier, Nuria Galiana and Yuval Zelnik for helpful discussions and review of previous versions of this manuscript. Our work has benefited greatly from the thorough and constructive reviews of Frédéric Barraquand, Kévin Cazelles, Kevin McCann, and an anonymous reviewer. This work was supported by the TULIP Laboratory of Excellence (ANR-10-LABX-41) and by the BIOSTASES Advanced Grant, funded by the European Research Council under the European Union’s Horizon 2020 research and innovation programme (grant agreement No 666971). This preprint has been reviewed and recommended by PCI Ecology (https://dx.doi.org/10.24072/pci.ecology.100017).

## Appendices

The Appendices are organized as follows: Appendix A through D provides a self-contained presentation of the mathematical foundations of our variability theory. Appendix E through H provide details concerning specific applications considered in the main text: two-species communities in Appendix E, complex Lotka-Volterra communities in appendices F and G, and the link between abundance statistics and variability in Appendix H. A list of the most important notation used in the Appendices is given in Table A1.

### Appendix A: Response to white-noise perturbation

We describe the response of a linear dynamical system, representing the dynamics of displacement of species around an equilibrium value, to a white-noise perturbation. Stochastic perturbations in continuous time are mathematically quite subtle (see, e.g., 37). However, in the setting of linear dynamical systems, the effect of a white-noise perturbation can be analyzed relatively easily. Because this analysis is not readily available in the ecology literature, we present here a short overview. We start from a fomulation in vector notation,

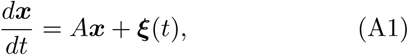

where ***x*** = (*x*_*i*_) denotes the vector of species displacements, ***ξ*** = (*ξ*_*i*_) the vector of species perturbations, and *A* = (*A*_*ij*_) the community matrix.

Suppose that the perturbation ***ξ*** (*t*) consists in a sequence of pulses. We denote the times at which these pulses occur by *t*_*k*_, and the corresponding pulse directions by **u**_k_ = (*u*_*k,i*_). The multi-pulse perturbation can then be written as

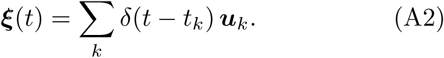

where we have used the Dirac delta function *δ* (*t*).

**TABLE A1:**
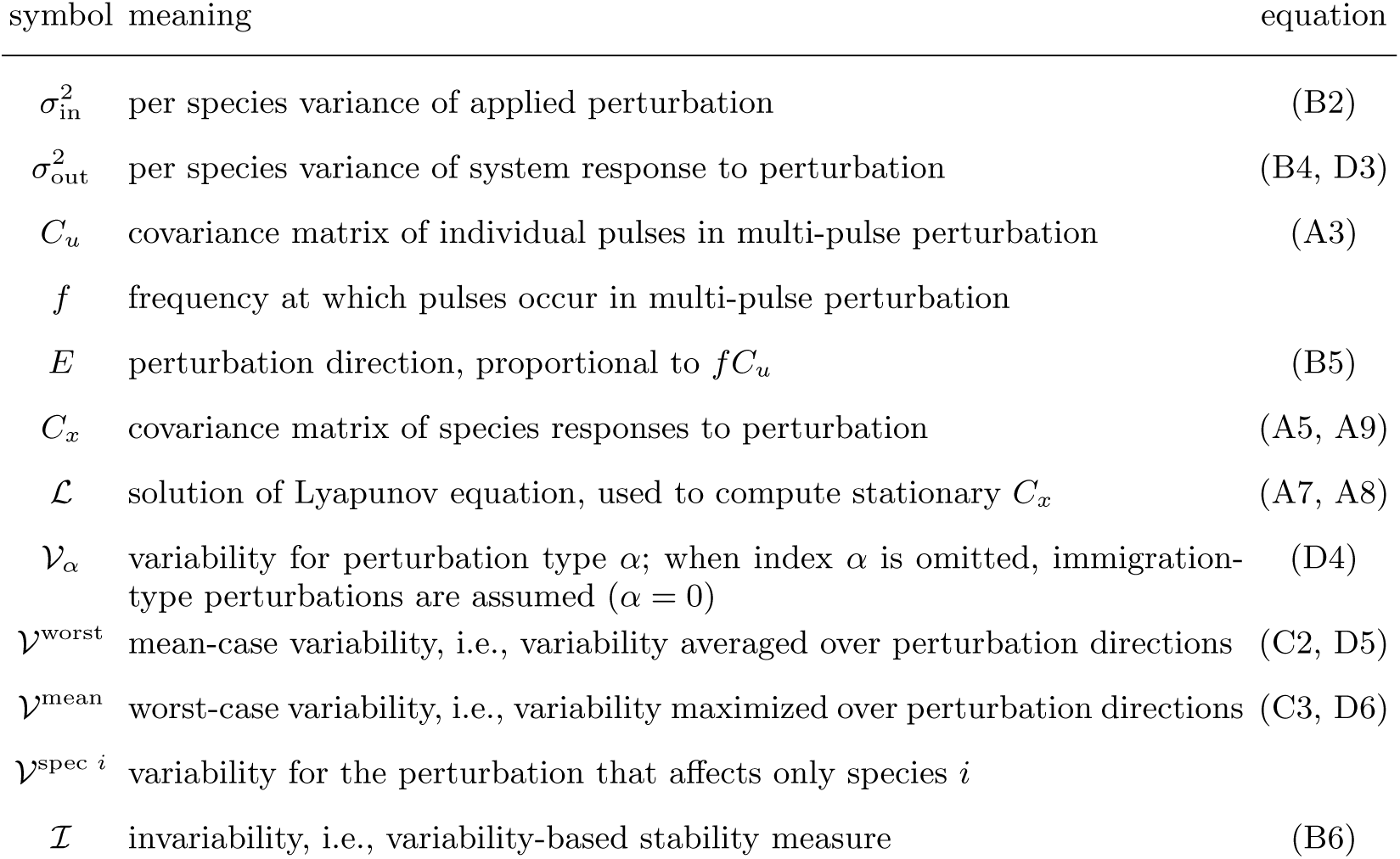
Notation used throughout the Appendices

We model both the pulse times *t*_*k*_ and the pulse directions ***u***_*k*_ as random variables. Specifically, we assume that the pulse times are distributed according to a Poisson point process with intensity *f*. This means that the probability that a pulse occurs in a small time interval of length. Δ*s* is equal to *f* Δ*s*, and that this occurrence is independent of any other model randomness. We denote the average over the pulse times *t*_*k*_ by 𝔼 _*f*_.

Furthermore, we assume that the pulse directions ***u***_*k*_ are independent (mutually independent, and independent of any other model randomness) and identically distributed. They have zero mean, and their second moments are given by the covariance matrix *C*_*u*_. That is, denoting the average over the pulse directions ***u***_*k*_ by 𝔼_*u*_, we have 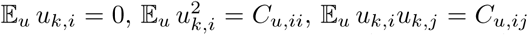 and 𝔼 _*u*_ *u*_*k,i*_ *u*_*𝓁,j*_ = 𝔼_*u*_ *u*_*k,i*_ *u*_,*𝓁, j*_ = 0 for *i* ≠ *j* and *k* ≠ *l*. The latter equations can be written in vector notation,

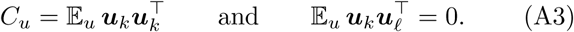

We use this information to compute the statistics of species displacements ***x***(*t*). Because the system response to a single pulse perturbation at time *t*_*k*_ in directon ***u***_*k*_ is equal to 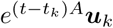, the system response to the sequence (A2) of pulse perturbations is equal to

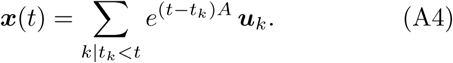

Taking the mean over the perturbation directions, we obtain

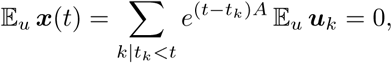

showing that the species displacements fluctuate around the unperturbed equilibrium.

Next, we compute the covariance matrix of the species displacements,

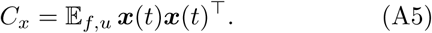

We substitute the response to the multi-pulse perturbation, eq. (A4),

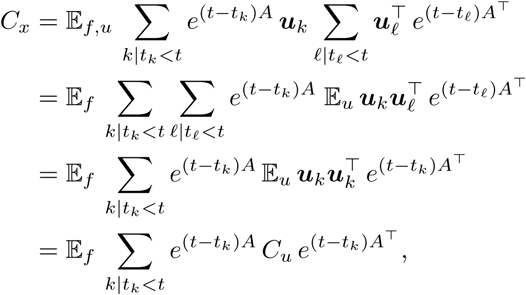

where we have used eq. (A3). To take the average over the pulse times, we partition the time axis in small intervals of length Δ*s*. Writing *s*_*n*_ = *n*Δ*s* for any integer *n*, we get

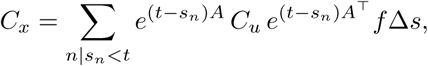

because the contribution of term *n* is equal to 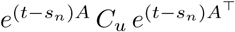 with probability f Δ*s*, and zero otherwise. Assuming that the time intervals. Δ*s* are in-finitesimal, we find the integral

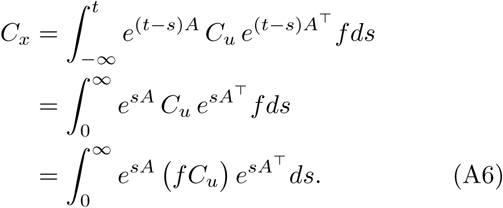

Hence, we have obtained the stationary covariance matrix of the species displacements under a stochastic multipulse perturbation.

A white-noise perturbation corresponds to a special case of the stochastic multi-pulse perturbation, namely, to the case of extremely frequent pulses (large f) of extremely small size (small ‖***u***‖). More precisely, we have to take the coupled limit *f* → ∞ and *C*_*u*_ → 0 while keeping *f C*_*u*_ constant. Because eq. (A6) depends on *f* and *C*_*u*_ through the product *fC*_*u*_ only, the same expression is also valid for white-noise perturbations.

Alternatively, the stationary covariance matrix *C*_*x*_ can be obtained by solving the so-called Lyapunov equation,

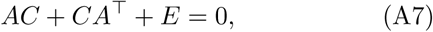

where *E* is the covariance matrix characterizing the white noise, equal to *f C*_*u*_ in our case. Indeed, it can be verified that eq. (A6) satisfies eq. (A7),

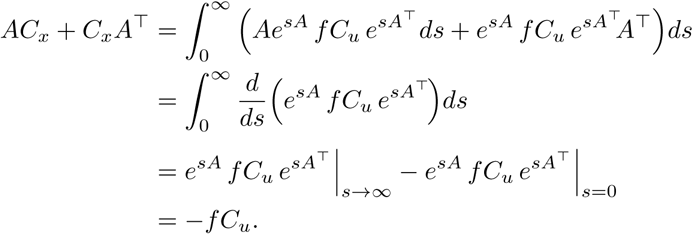

For a stable matrix *A* this is the unique solution of the Lyapunov equation, for which we introduce the short-hand notation *ℒ* (*A, E*),

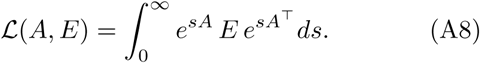

Hence, we can write

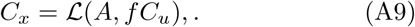

From a numerical viewpoint, the covariance matrix *C*_*x*_ can be easily obtained by solving the Lyapunov eq. (A7), which can be written as a system of *S*^2^ linear equations, rather than by computing the integral in (A8). Note also that solution of Lyapunov equation is linear in the perturbation covariance matrix,

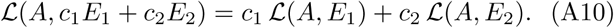

### Appendix B: Construction of variability measure

We explain the construction of the variability measure 𝒱, see eq. (4) in the main text. The construction is based on the comparison of the intensity of the system response relative to the intensity of the applied perturbation. It should be stressed that, while we take special care of quantifying these intensities in a reasonable way, alternative choices are possible.

#### a. Perturbation intensity

A reasonable measure of the perturbation intensity should increase with the number of pulses and the intensity of each pulse separately. In particular, we expect it to be proportional to the pulse frequency f and to some function of the pulse covariance matrix *C*_*u*_.

We propose to look at the squared displacements ‖***u***_*k*_ ‖^2^ induced by pulses ***u***_*k*_. The accumulated squared displacement in time interval [*t, t* + *T*] is

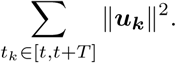

Taking the average over pulse times and pulse directions,

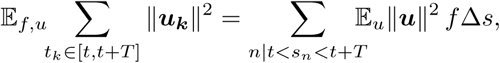

where we have partitioned the time axis in small intervals of length. Δ*s* (see derivation of eq. (A6)). Then,

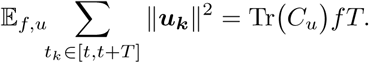

The result is proportional to the length *T* of the considered time interval. The average accumulated squared displacement per unit of time is

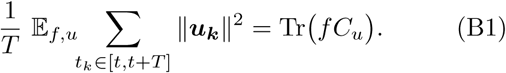

As expected, this quantity is proportional to the pulse frequency f and increases with the pulse covariance matrix *C*_*u*_. Note also that *f* and *C*_*u*_ appear as a product, so that the expression is compatible with the white-noise limit.

Eq. (B1) quantifies the intensity of the perturbation applied to the entire ecosystem. This measure is not directly appropriate to normalize the pertubation intensity across systems. Indeed, when keeping the total perturbation intensity constant, the perturbation applied to a given species would be weaker in a community with a larger number of species. To eliminate this artefact, we normalize the perturbation intensity on a per species basis. Thus, we propose to quantify the perturbation intensity as

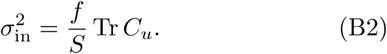

#### b. Response intensity

We measure the intensity of the system response in terms of the covariance matrix *C_x_*. This matrix encodes the statistical properties of the abundance (or biomass) fluctuations in stationary state. For example, species abundance *x*_*i*_ (*t*) fluctuates around its equilibrium value *N*_*i*_ with variance *C*_*x,ii*_. More generally, we can describe the fluctuations of any function φ of species abundance. The dynamics near equilibrium are

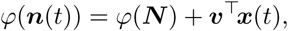

where vector ***υ*** = ∇*φ* is the gradient of the function *φ* evaluated at the equilibrium ***N.*** This vector gives the direction in which the system fluctuations are observed. Then, denoting the temporal mean and variance by 𝔼_*t*_ and Var_*t*_, we have

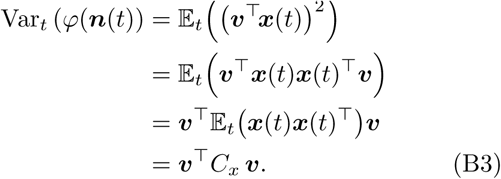

We use this variance to quantify the intensity of the system response. Rather than choosing a particular vector ***υ***, we consider the average over all observation directions. Specifically, we restrict attention to unit vectors ***υ*** and average over the uniform distribution of such vectors. Denoting this average by 𝔼*_v_*, we get

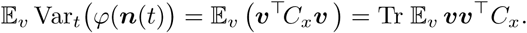

It follows from species symmetry that the average 𝔼_v_ ***υ υ***^T^ is proportional to the unit matrix. Moreover, because Tr ***υ υ***^*T*^ = 1 for all vectors ***v***, the constant of proportionality is equal to 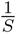. Hence,

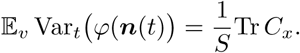

Therefore, we propose to quantify the response intensity as

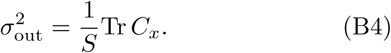

#### c. Variability and invariability

We define variability 𝒱 as the ratio of the response intensity 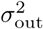 and the perturbation intensity 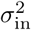,

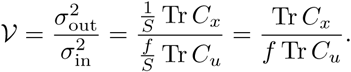

Substituting eq. (A9) for *C*_*x*_, we get

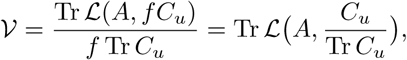

where we have used the linearity property (A10). We see that only the normalized perturbation covariance matrix matters in this expression. That is, the variability measure focuses on the directional effect of the perturbation. We make this dependence explicit in the notation, and write

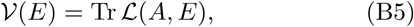

where 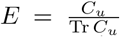 is the perturbation direction, i.e., a covariance matrix with unit trace.

Variability is inversely related to stability: the more variable an ecosytem, the less stable it is. For purpose of comparison, we construct a stability measure based on variability 𝒱 (*E*), which we call invariability ℐ (*E*),

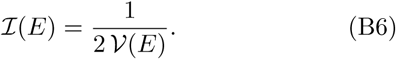

The factor 2 in this definition guarantees that we recover asymptotic resilience for the simplest dynamical systems. To see this, consider a system of *S* non-interacting species, in which all species have the same return rate λ. The community matrix is equal to *A* = –λ𝕀 where 𝕀 denotes the identity matrix. From the Lyapunov equation (A7) we get the stationary covariance matrix 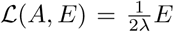. Therefore, 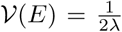 and ℐ (*E*) = λ, which is equal to the asymptotic resilience of this example system.

### Appendix C: Worst-case and mean-case variability

**Worst-case variability** is defined as

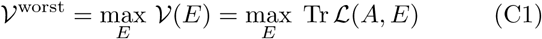

where the maximum is taken over perturbation directions, i.e., over covariance matrices *E* with Tr *E* = 1. The function Tr ℒ (*A, E*) is linear in the perturbation direction *E*, see eq. (A10), and the set of perturbation directions is convex. Hence, the maximum is reached at an extreme point, that is, on the boundary of the set. The extreme points are the purely directional perturbations (see Appendix E for the argument in the two-species case), so that the maximum is reached at a purely directional perturbation. Arnoldi et al. [4] showed that the worst-case variability can be easily computed, namely, as a specific norm of the operator *Â^-1^* that maps *E* to ℒ (*A, E*). Concretely, defining *Â* = *Â* ⨂ 𝕀 + 𝕀 ⨂ *A*,

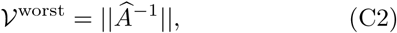

where ∥ · ∥ stands for the spectral norm of *S*^2^ × *S*^2^ matrices.

To define **mean-case variability** 𝒱^mean^, we assume a probability distribution over the perturbation directions, and compute the mean system response over this distribution. Due to the linearity property (A10), this mean intrinsic perturbation. The derivation leading to eq. (B2) is still valid. However, to compute the covariance matrix of the species displacements, we use the covariance matrix of the pulses in the total perturbation. This correresponse is equal to the response to the mean perturbation direction. Hence, we do not have to specify the full probability distribution over the perturbation directions; it suffices to determine the mean perturbation direction. As can be directly verified in the two-species case (Appendix E), if, averaged over the distribution of perturbation directions, perturbation intensities are evenly distributed across species, and positive and negative correlations between species perturbations cancel out, then the mean perturbation direction is adirectional. This corresponds to the center of the set of perturbation directions (in the two-species case the disc center represented in Fig. 3), and is proportional to the identity matrix, that is, 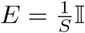. Therefore,

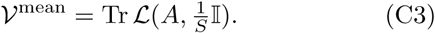

### Appendix D: Perturbation types and variability

The perturbation type (environmental-, demographic-or immigration-type) affects how the perturbation intensity is distributed across species. Therefore, it also affects our measure of variability, as defined in Appendix B. Here we describe how the variability definition has to be modified.

We defined variability measure (B5) as the intensity of the system response relative to the intensity of the applied perturbation. To quantify the perturbation intensity in the case of abundance-dependent perturbations, we distinguish the intrinsic effect of the perturbation on a species, which does not depend on the species’ abundance, and the total effect of the perturbation on the species, which does depend on abundance. We propose to express the perturbation intensity in terms of the intrinsic perturbation, while it is the total perturbation that acts on the species dynamics.

Formally, for species *i*, we denote the intrinsic perturbation by 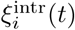 and the total perturbation by 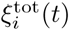. Then, for a type-α perturbation, we have

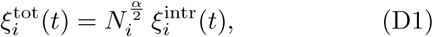

where *N*_*i*_ is the abundance of species *i*. Thus, the intrinsic perturbation ξ^intr^ (*t*) can be interpreted as the per capita perturbation strength. Eq. (D1) can be written in vector notation as

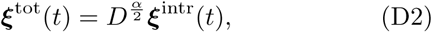

where *D* is the diagonal matrix whose entries are species equilibrium values (*D*_*ii*_ = *N*_*i*_).

Both the intrinsic and total perturbation are multipulse. If we denote the pulses of the intrinsic perturbation by ***u****_k_*, then, by eq. (D2), those of the total perturbation are 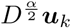. Then, to quantify the perturbation intensity, we use the covariance matrix of the pulses in the intrinsic perturbation. The derivation leading to eq. (B2) is still valid. However, to compute the covariance matrix of the species displacements, we use the covariance matrix of the pulses in the total perturbation. This corresponds sponds to replacing *C*_*u*_ by 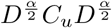 in the derivation of eq. (B4), so that we get

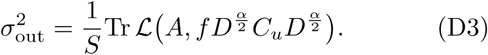

The variability measure for a type-α perturbation becomes

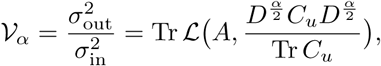

or, in terms of the (intrinsic) perturbation direction *E*,

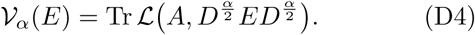

Applying the same arguments as in Appendix C, we find that worst-case variability,

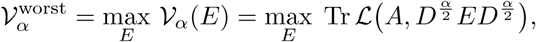

is attained at a perfectly correlated perturbation. If we define the operator (an *S*^2^ × *S*^2^ matrix)

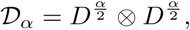

then the worst case-variability can be computed as

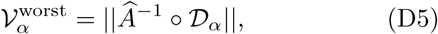

Where ‖ · ‖ is the spectral norm for *S*^2^ × *S*^2^ matrices. On the other hand, the mean-case variability,

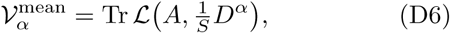

is attained by the uniform, uncorrelated perturbation.

### Appendix E: Perturbation directions in two dimensions

Variability spectra are built on the notion of perturbation directions. They are characterized by a covariance matrix *E* with Tr *E* = 1. To gain some intuition, we study the set of perturbation directions in the case of two species.

Any perturbation direction *E* in two dimensions can be written as

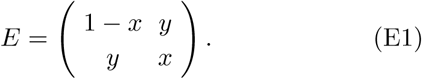

with 0 ≤ *x* ≤ 1 and *y*^2^ ≤ *x*(1 – *x*). The first inequality guarantees that the elements on the diagonal are variances, i.e., positive numbers. The second inequality guarantees that the off-diagonal element is a proper covariance, in particular, that the correlation coefficient is contained between –1 and 1. Note that matrix (E1) has always Tr *E* = 1.

It follows from eq. (E1) that the set of perturbation directions in two dimensions is parameterized by two numbers *x* and *y*. Using these numbers as axes of a two-dimensional plot, we see that the set of perturbation directions corresponds to a disc with radius 0.5 and centered at (0.5, 0) (see Fig. 3).

It is instructive to study the position of specific perturbation directions on the disc. The point (0, 0) corresponds to a perturbation affecting only the first species, whereas point (1, 0) is a perturbation only affecting the second species. More generally, any point on the boundary of the disc correspond to a multi-pulse perturbation for which the individual pulses have a fixed direction. For example, the point (0.5, 0.5) is a perturbation for which each pulse has the same effect on species 1 and species 2, whereas the perturbation corresponding to point (0.5, 0.5) consists of pulses that affect the two species equally strongly, but in an opposite way. Perturbations on the boundary are *perfectly correlated*.

The perturbations towards the center of the disc are composed of pulses with more variable directions. For example, a multi-pulse perturbation for which half of the pulses affect only the second species, and the other pulses affect the two species equally strongly corresponds to the point 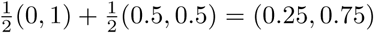. The mixture of different pulse directions is the strongest at the center of the disc (0.5, 0). Examples of ways to realize this perturbation are 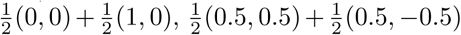 and 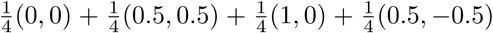. In each of these example, the pulses have their intensities, averaged over time, evenly distributed across species, and affect them, again averaged over time, in an uncorrelated way. The perturbation corresponding to the point (0.5, 0) is thus evenly distributed across species but uncorrelated in time.

### Appendix F: Random Lotka-Volterra model

The communities used in Figs. 4, 5 and 6 are constructed from the Lotka-Volterra model with random parameters. We consider a pool of species governed by the dynamics

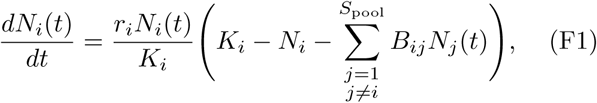

and we let the dynamics settle to an equilibrium community of *S* remaining species. By drawing random values for the parameters – growth rates r_i_, carrying capacities *K*_i_, and competition coefficients *B*_*ij*_ – we generate communities of various diversity.

For the communities in Fig. 4, we set *S*_pool_ = 50, and chose the parameter values as follows,

*r*_*i*_ randomly drawn from 𝒩 (1, 0.2), a normal distribution with mean 1 and standard deviation 0.2 (independent draws for different species)

*K*_*i*_ drawn from 𝒩 (1, 0.2)

*B*_*ij*_ half of the competition coefficients are set equal to 0; the other half are drawn from 𝒩 (0.1, 0.1).

This procedure resulted in a community of *S* = 40 persistent species. Note that some of the competition coefficients can be negative, so that there can be positive interactions (e.g. facilitation).

For the communities in the top row of Fig. 5, we followed the same procedure, except that we changed the way of generating the competition coefficients *B*_*ij*_. In the case without interactions, all *B*_*ij*_ were set zero; in the case with weak interactions, the non-zero coefficients *B*_*ij*_ were drawn from 𝒩 (0.02, 0.02); and in the case with strong interactions, the non-zero *B*_*ij*_ were drawn from 𝒩 (0.1, 0.1), as for the community of Fig. 4.

We applied a similar procedure to obtain the bottom row of Fig. 5, but for these communities the growth rates *r*_*i*_ and carrying capacities *K*_*i*_ were not drawn independently. Instead, we first drew auxiliary variables *a*_*i*_ from 𝒩 (1, 0.2), *b*_*i*_ from 𝒩 (1, 0.1) and *c*_*i*_ from 𝒩 (1, 0.1), and then set *r*_*i*_ = *b*_*i*_*a*_*i*_ and *K*_*i*_ = *c*_*i*_/*a*_*i*_.

For the communities of Fig. 6, we varied the size of the species pool *S*_pool_ so that the realized species richness covered the range from 1 to 20. Specifically, we drew *S*_pool_ randomly from 1 to 100, and generated the parameter values as in Fig. 4. We repeated this procedure many times, until obtaining 1000 communities for each value of realized species richness S from 1 to 20. Then, for each realized community, and for each of the three perturbation types (α = 0, α = 1 and α = 2), we generated 1000 random perturbations leading to a variability distribution of 1000 values. From the variability distributions we extracted median, 5th and 95th percentile, and minimum and maximum. For the realized communities we computed asymptotic resilience, worst-case variability and the prediction for the median. Finally, we computed the median of these statistics and predictions, all represented in Fig. 6.

### Appendix G: Genericity in strongly interacting communities

We give some elements as to why the behavior reported in Figs. 4 and 5 in the main text can be expected to be a general trend in diverse communities of interacting species. Denote by 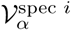 the community variability induced by a type-α perturbation that is fully focused on a single species *i*. We are interested in the relationship between this variability and the equilibrium abundance *N*_*i*_ of the perturbed species *i*.

**FIG. G1:**
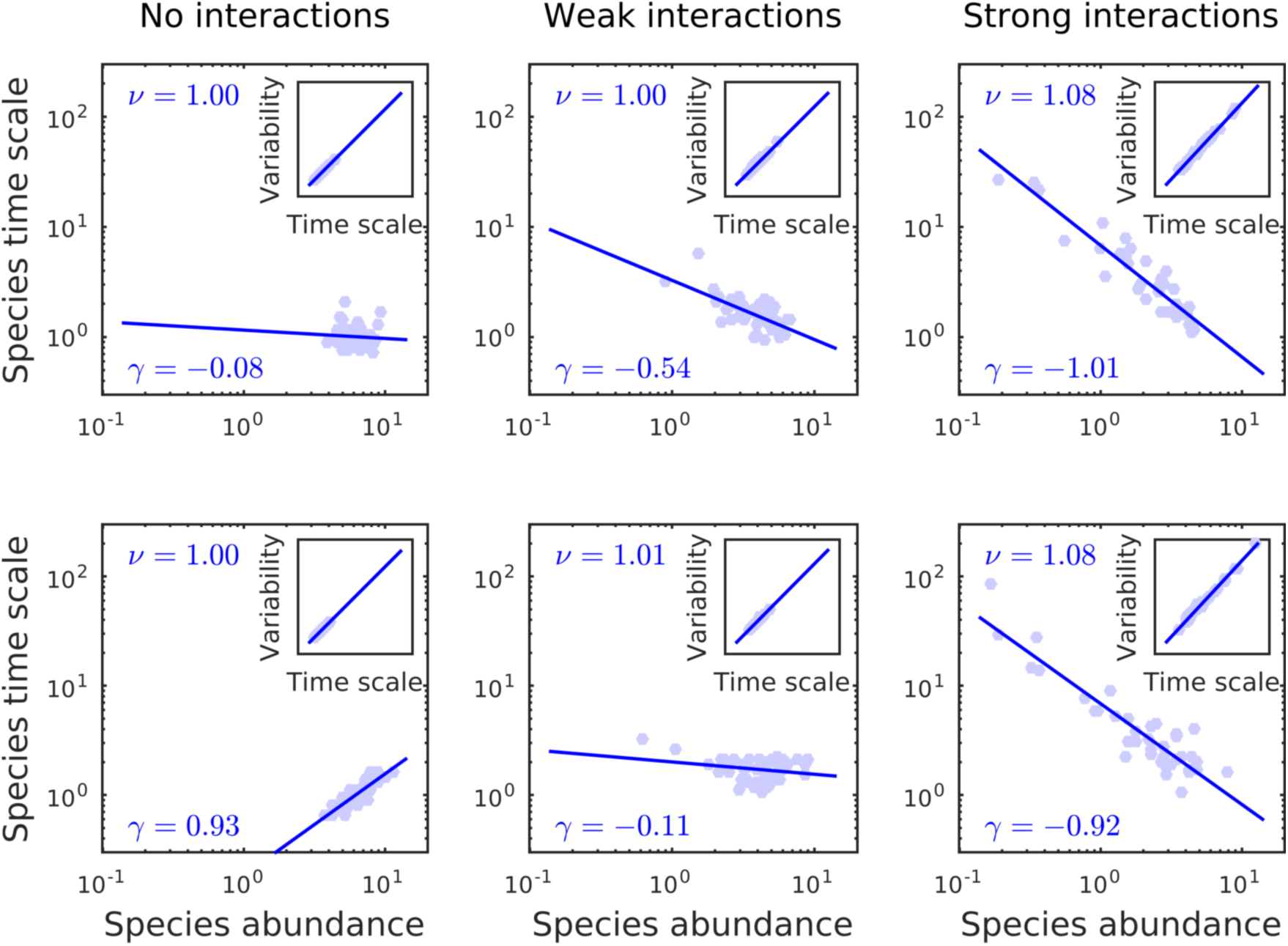
Clarifying the relationship between abundance of perturbed species and community variability. In Appendix G we introduce the auxiliary variable *τ*_*i*_, the characteristic time scale of species *i*, to explain the relationship between variability 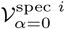 and abundance *N*_*i*_. For the six communities of Fig. 5 in the main text, we plot *τ*_*i*_ vs *N*_*i*_ in the main panels, and 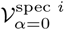 vs *τ*_*i*_ in the inset panels. We fit a power law to each of these relationships, using linear regression on the log-log plot. The estimated exponents γ (for the data *τ*_*i*_ vs *N*_*i*_) and γ (for the data 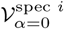 vs *τ*_*i*_) are reported in the panels.

First, note that for single-species perturbations the variability metrics 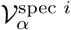 for different perturbation types *α* are directly linked. From definition (D4) we get that

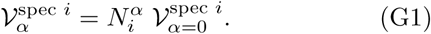

Hence, it suffices to study the behavior of 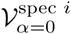.

Next, consider again the Lotka-Volterra dynamics (F1) from the perspective of a focal species *i*. If a stable equilibrium exists in which the focal species survives, small displacements from equilibrium *x*_*i*_ = *N*_*i*_(*t*) - *N*_*i*_ are met with the dynamics

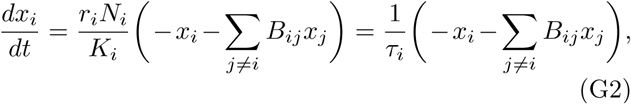

where 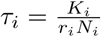 has units of time. We claim that *τ*_*i*_ sets a characteristic time scale of the focal species dynamics; it measures the typical time it takes for the species to recover from a perturbation that displaces it from its equilibrium. This species response time is directly related to the species’ variability 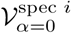: the slower the species, the larger the impact of a repeated perturbation acting on this species, and the larger the induced variability.

We illustrate the relationship between *τ*_*i*_ and 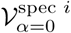 in Fig. G1 (inset panels). For the six communities of Fig. 5, we fit the power-law relationship

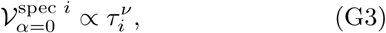

where the index *i* runs over the set of persistent species. The estimates of the exponent *v* (using linear regression on the log-log plot) are all close to one. This result is obvious for the communities without interactions, for which 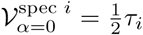 (left-hand panels). But the same result remains valid in the presence of interactions. We find that interactions do not substantially modify the time scale on which a species responds to perturbations affecting only that species.

Therefore, to study the relationship between *N*_*i*_ and 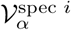, we can restrict to the simpler relationship - between *N*_*i*_ and 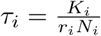, which is determined by the correlations between growth rates *r*_*i*_, carrying capacities *K*_*i*_ and equilibrium abundances *N*_*i*_. Fig. G1 (main panels) shows this relationship for the six communities of Fig. 5. Fitting the power law

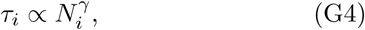

we find various estimates for the exponent γ. Without interactions, we have *N*_*i*_ = *K*_*i*_, and hence, 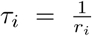. If growth rates and carrying capacities are drawn independently, abundance and response time are unrelated, leading to γ ≈ 0 (Fig. G1, upper-left panel). Alternatively, if growth rates and carrying capacities satisfy some trade-off, higher abundance (larger *K*_*i*_) is associ-ated with longer response time (smaller *r*_*i*_), leading to γ > 0 (Fig. G1, lower-left panel). When increasing the interactions, the link between *N*_*i*_ and *K*_*i*_ becomes weaker. Indeed, from the equilibrium condition for species *i* we have

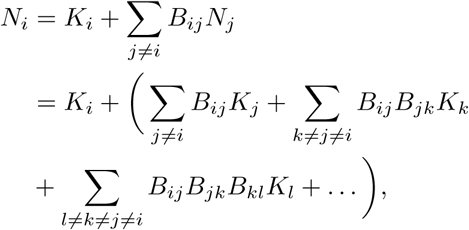

where in the second line we have used the equilibrium condition for the other species. For sufficiently strong interactions, the terms between brackets dominate the term *K*_*i*_, so that *N*_*i*_ and *K*_*i*_ become unrelated. In this case, we have 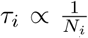, leading to γ ≈-1: more abundant species have faster dynamics and smaller response time. This limiting case is observed both if *r*_*i*_ and *K*_*i*_ are independent, and if they satisfy a trade-off (Fig. G1, right-hand panels).

Finally, putting together eqs. (G1, G3, G4), we get

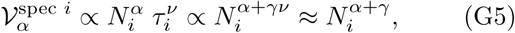

where in the last step we have used that *v* ≈ 1. The relationship between abundance of perturbed species and community variability is strongly determined by the exponent γ, that is, by the relationship between abundance *N*_*i*_ and response time *τ*_*i*_. In the case of weak interactions, the latter relationship depends on the assumed link between growth rate *r*_*i*_ and carrying capacity *K*_*i*_, so that no unambiguous relationship is to be expected be-tween abundance and variability. However, in the limit of strong interactions, we have γ ≈ −1 and

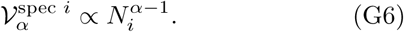

Hence, for immigration-type perturbations (*α* = 0) variability is inversely proportional to the abundance of the perturbed species. In contrast, for environmental perturbations (*α* = 2), variability is directly proportional to the abundance of the perturbed species. These are the relationships depicted in Figs. 4 and 5 of the main text.

### Appendix H: Variability and abundance statistics

From the observed relationship between abundance and variability (Figs. 4 and 5), patterns for worst-and mean-case variability can be deduced. This reveals a connection between stability and diversity metrics.

Denote by 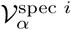 the community variability induced by a type-*α* perturbation fully focused on species *i*. We start from the power-relationship (G6), linking this variability and the equilibrium abundance of species *i*. As argued in Appendix G, we expect this relationship to hold for sufficiently strong interactions.

For immigration-type perturbations (*α* = 0), worst-case variability is approached by taking the maximum over species which gives

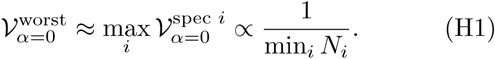

so that the worst case is governed by the rarest species. Because the abundance of the rarest species typically decreases with diversity, the corresponding diversity-stability relationship is decreasing. For mean-case variability, averaging over species individual contributions, we get

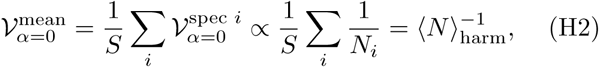

where ⟨*N*⟩ _harm_ stands for the harmonic mean of species abundances. Mean abundance typically decreases with diversity, so that the corresponding diversity-stability relationship is decreasing.

When caused by environmental-type perturbations (*α* = 2), worst-case variability is approached by taking the maximum over species, giving

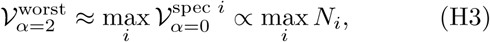

so that the worst case is governed by the most abundant species. For mean-case variability we get

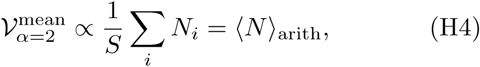

the arithmetic mean of species abundances. Mean abundance typically decreases with diversity, so that the corresponding diversity-stability relationship is increasing.

Note that when caused by demographic-type perturbations (*α* = 1) the species-by-species approach does not work: demographic variability probes a collective property of the community. The different relationships between abundance and variability are illustrated in Fig. H1.

**FIG. H1:**
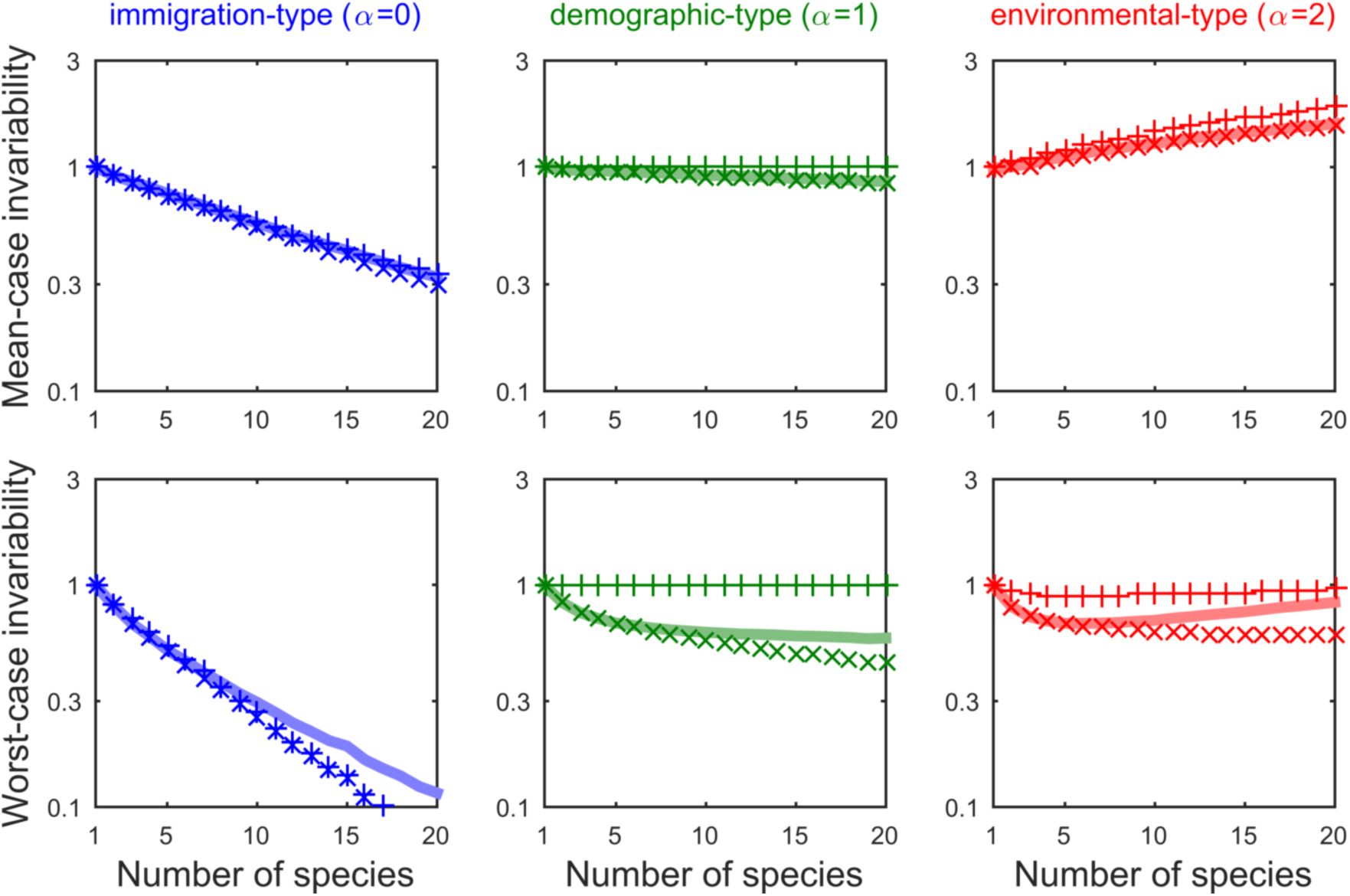
Invariability and species abundance. Top row: mean-case, bottom row: worst-case. ×-marks: analytical formula; +-marks: approximation in terms of abundance (see Appendix H); thick line: simulation results. For immigration-type perturbations (first column, in blue), mean-case invariability scales as the harmonic mean abundance (see eq. (H2)), which decreases with diversity. Worst-case invariability scales as the abundance of the rarest species. On the other hand, in response to environmental-type perturbations (third column, in red), mean-case variability scales as the arithmetic mean abundance (see eq. (H4)) so that invariability increases. Worst-case variability scales as the abundance of the most common species. In between (second column, in green), for demographic-type perturbations, neither worst-nor mean-case invariability is determined by statistics of species abundances.

