## Supplementary material for "The inherent multidimensionality of temporal variability: How common and rare species shape stability patterns": Matlab code used to generate figures: code_fig3.m

```
%%%%%%%%%%%%%%%%%%%%%%%%%%%%%%%%%%%%%%%%%%%%%%%%%%%%%%%%%%%%%%%%%%%%%%%%%%%
% This is Matlab code for the results presented in Figure 3 of
% Arnoldi et al. The variability spectrum of ecological communities:
% How common and rare species shape stability patterns
%%%%%%%%%%%%%%%%%%%%%%%%%%%%%%%%%%%%%%%%%%%%%%%%%%%%%%%%%%%%%%%%%%%%%%%%%%%

% we compute the variability measure (equation (5) of the paper) for all
% possible perturbation covariance matrices, and for three perturbation
% types. the perturbation covariance matrix is given by C=[1-x y;y x]. The
% perturbation types are al=0 (panel A), al=1 (panel B) and al=2 (panel C).

N=[.4 3]; % equilibrium abundances
A=[-1 .1;-4 -1]; % community matrix
xs=linspace(0,1); % range of x-values in covariance matrix
ys=linspace(-.5,.5); % range of y-values in covariance matrix
als=[0 1 2]; % set of perturbation types
res=zeros(3,length(xs),length(ys));
    % variable to store variability results
for ix=1:length(xs),
    for iy=1:length(ys),
        x=xs(ix);
        y=ys(iy);
        if y >= -sqrt(x*(1-x)) && y <= sqrt(x*(1-x)),
            % construct peturbation covariance matrix
            E=[1-x y;y x];
            % compute variability measure
            res(:,ix,iy)=variability_perturbation(N,A,E,[0 1 2]);
        else
            % parameters x,y do not correspond to a covariance matrix
            res(:,ix,iy)=nan;
        end
    end
end
```
