## Supplementary material for "The inherent multidimensionality of temporal variability: How common and rare species shape stability patterns": Matlab code used to generate figures: code_fig6.m

```
%%%%%%%%%%%%%%%%%%%%%%%%%%%%%%%%%%%%%%%%%%%%%%%%%%%%%%%%%%%%%%%%%%%%%%%%%%%
% This is Matlab code for the results presented in Figure 6 of
% Arnoldi et al. The variability spectrum of ecological communities:
% How common and rare species shape stability patterns
%%%%%%%%%%%%%%%%%%%%%%%%%%%%%%%%%%%%%%%%%%%%%%%%%%%%%%%%%%%%%%%%%%%%%%%%%%%

%%% FIRST STEP: GENERATE COMMUNITIES %%%

% we generate a database of communities. the database contains 20000
% communities in total. the number of persisting species ranges from 1 to
% 20. there are 1000 communities for each number of species.

% these communities are obtained by repeating the following procedure:
% (1) generate the size of the species pool, by drawing a random number
%   between 1 and 100 (uniform on log-scale)
% (2) generate the parameters of the species pool, by drawing randomly
% . intrinsic growth rates: normal with mean 1 and standard deviation 0.2
% . carrying capacities: normal with mean 1 and standard deviation 0.2
% . intraspecific interaction strength: fixed, equal to 1
% . interspecific interaction strengths: equal to 0 with probability 0.5,
%   otherwise drawn from a normal distribution with mean 0.1 and standard
%   deviation 0.1
% (3) simulate the Lotka-Volterra model for a long simulation time (1e5
%   time units), starting from low species abundances
% (4) check whether the final state is a stable equilibrium, and whether
%   the number of persisting species lies between 1 and 20.

comms=struct('A',cell(1000,20), ...
             'B',cell(1000,20), ...
             'N',cell(1000,20));
         % structure to store communities
ncomms=zeros(1,20); % number of communities in database

while any(ncomms<1000),
    % (1) generate size of species pool
    spool=round(100^rand(1));

    % (2) generate parameters of species pool
    rr=1+.2*randn(spool,1); % intrinsic growth rates
    kk=1+.2*randn(spool,1); % carrying capacities
    bb=.1+.1*randn(spool);
    bb=bb.*(rand(spool)<.5);
    bb=bb-diag(diag(bb))+eye(spool); % interaction matrix

    % (3) simulate Lotka-Volterra model
    A=rr;
    B=-diag(rr./kk)*bb;
    x0=1e-3*ones(spool,1);
    opt1=odeset('abstol',1e-8,'reltol',1e-8,'nonnegative',1:spool, ...
        'jacobian',@(t,x) diag(A+B*x)+diag(x)*B);
    [~,Y]=ode15s(@(t,x) x.*(A+B*x),[0 1e5],x0,opt1);
    N=Y(end,:);
    N=N(:); % species abundances at end of simulation
    icomm=find(N>1e-6); % indices of persistent species
    scomm=length(icomm); % number of persistent species
    Br=B(icomm,icomm); 
    Ar=A(icomm);
    Nr=N(icomm);
    opt2=optimset('display','off','tolfun',1e-8,'tolx',1e-8);
    Nr=fsolve(@(x) x.*(Ar+Br*x),Nr,opt2);

    % (4) check appropriateness equilibrium
    isequi=(norm(Nr.*(Ar+Br*Nr))<1e-6);
    isfeas=all(Nr>1e-6); % feasibility check
    jacr=diag(Ar+Br*Nr)+diag(Nr)*Br;
    isstab=(max(real(eig(jacr)))<0); % stability check
    if isequi && isfeas && isstab && scomm<=20 && ncomms(scomm)<1000,
        ncomms(scomm)=ncomms(scomm)+1;
        comm=struct('A',Ar,'B',Br,'N',Nr);
        comms(ncomms(scomm),scomm)=comm;
    end
end

%%% SECOND STEP: COMPUTE STABILITY MEASURES %%%

% for each randomly generated communities we compute the following set of
% stability measures:
% . asyresil: asymptotic resilience
% . varmean: mean-case variability, for 3 perturbation types alpha=0,1,2
%   (size 1x3)
% . varworst: worst-case variability, for 3 perturbation types alpha=0,1,2
%   (size 1x3)
% . varsimul: variability statistics for random perturbation directions;
%   we generate 1000 random perturbation directions, and compute 5 summary
%   statistics (minimum, 5% percentile, median, 95% percentile, maximum),
%   for 3 perturbation types alpha=0,1,2 (size 5x3)

stabs=struct('asyresil',cell(1000,20), ...
             'varmean',cell(1000,20), ...
             'varworst',cell(1000,20), ...
             'varsimul',cell(1000,20));
         % structure to store stability values

for nsp=1:20,
    for icomm=1:1000,
        A=comms(icomm,nsp).A;
        B=comms(icomm,nsp).B;
        N=comms(icomm,nsp).N;
        J=diag(A+B*N)+diag(N)*B;
        asyresil=-max(real(eig(J)));
        varmean=variability_meancase(N,J,[0 1 2]);
        varworst=variability_worstcase(N,J,[0 1 2]);
        varperts=zeros(1000,3);
        for ipert=1:1000,
            pert=rand(nsp,1);
            pert=pert/norm(pert);
            E=pert*pert';
            varperts(ipert,:)=variability_perturbation(N,J,E,[0 1 2]);
        end
        varsimul(1,:)=min(varperts);
        varsimul(2:4,:)=prctile(varperts,[5 50 95]);
        varsimul(5,:)=max(varperts);
        stab=struct('asyresil',asyresil,'varmean',varmean, ...
            'varworst',varworst,'varsimul',varsimul);
        stabs(icomm,nsp)=stab;
    end
end
```
