## Supplementary material for "The inherent multidimensionality of temporal variability: How common and rare species shape stability patterns": Matlab code used to generate figures: code_fig45.m

```
%%%%%%%%%%%%%%%%%%%%%%%%%%%%%%%%%%%%%%%%%%%%%%%%%%%%%%%%%%%%%%%%%%%%%%%%%%%
% This is Matlab code for the results presented in Figures 4-5 of
% Arnoldi et al. The variability spectrum of ecological communities:
% How common and rare species shape stability patterns
%%%%%%%%%%%%%%%%%%%%%%%%%%%%%%%%%%%%%%%%%%%%%%%%%%%%%%%%%%%%%%%%%%%%%%%%%%%

%%% FIRST STEP: GENERATE COMMUNITIES %%%

% we generate communities by simulating the assembly process for a species
% pool of 50 species. species dynamics are governed by the Lotka-Volterra
% model with randomly drawn parameters.
% we consider 6 = 2*3 communities by combining
% . intrinsic growth rates and carrying capacities of the species in the
%   species pool are drawn independently
% . intrinsic growth rates and carrying capacities are drawn jointly (i.e.
%   not independently) such that they tend to be inversely proportional
% and
% . species in the species pool do not interact at all
% . species interact weakly
% . species interact strongly

% specifically,
% . in the case of independent r and K,
%   intrinsic growth rates are drawn from a normal distribution with mean
%   1 and standard deviation 0.2. carrying capacities are drawn from a
%   normal distribution with mean 1 and standard deviation 0.2
% . in the case of dependent r and K,
%   we first draw for each species an auxiliary variable from a normal
%   distribution with mean 1 and standard deviation 0.2. intrinsic growth
%   rates are obtained by multiplying the auxiliary variable by a normal
%   random variable with mean 1 and standard deviation 0.1. carrying
%   capacities are obtained by multiplying the reciprocal auxiliary
%   variable by a normal variable with mean 1 and standard deviation 0.1
% . in the case of no species interactions,
%   intraspecific interactions are fixed to 1 
%   interspecific interactions are fixed to 0
% . in the case of weak species interactions,
%   intraspecific interactions are fixed to 1 
%   interspecific interactions are fixed to 0 with probability 0.5, or
%   otherwise are drawn from a normal distribution with mean 0.02 and
%   standard deviation 0.02
% . in the case of strong species interactions,
%   intraspecific interactions are fixed to 1 
%   interspecific interactions are fixed to 0 with probability 0.5, or
%   otherwise are drawn from a normal distribution with mean 0.1 and
%   standard deviation 0.1

comms=struct('A',cell(2,3), ...
             'B',cell(2,3), ...
             'N',cell(2,3));
    % structure to store communities
spool=50; % size of species pool

for ic1=1:2,
    % ic1==1: r and K independent
    % ic1==2: r and K dependent (inversely proportional)
    for ic2=1:3,
        % ic2==1: no species interactions
        % ic2==2: weak species interactions
        % ic2==3: strong species interactions

        chkcomm=0;
        while ~chkcomm,
            % generate parameters of species pool
            if ic1==1,
                rr=1+.2*randn(spool,1); % intrinsic growth rates
                kk=1+.2*randn(spool,1); % carrying capacities
            elseif ic1==2,
                aux=1+.2*randn(spool,1);
                rr=aux.*(1+.1*randn(spool,1)); % intrinsic growth rates
                kk=1./aux.*(1+.1*randn(spool,1)); % carrying capacities
            end
            if ic2==1,
                bb=eye(spool); % interaction matrix
            elseif ic2==2,
                bb=.02+.02*randn(spool);
                bb=bb.*(rand(spool)<.5);
                bb=bb-diag(diag(bb))+eye(spool); % interaction matrix
            elseif ic2==3,
                bb=.1+.1*randn(spool);
                bb=bb.*(rand(spool)<.5);
                bb=bb-diag(diag(bb))+eye(spool); % interaction matrix
            end

            % simulate Lotka-Volterra model
            A=rr;
            B=-diag(rr./kk)*bb;
            x0=1e-3*ones(spool,1);
            opt1=odeset('abstol',1e-8,'reltol',1e-8,'nonnegative',1:spool, ...
                'jacobian',@(t,x) diag(A+B*x)+diag(x)*B);
            [~,Y]=ode15s(@(t,x) x.*(A+B*x),[0 1e5],x0,opt1);
            N=Y(end,:);
            N=N(:); % species abundances at end of simulation
            icomm=find(N>1e-6); % indices of persistent species
            scomm=length(icomm); % number of persistent species
            Br=B(icomm,icomm); 
            Ar=A(icomm);
            Nr=N(icomm);
            opt2=optimset('display','off','tolfun',1e-8,'tolx',1e-8);
            Nr=fsolve(@(x) x.*(Ar+Br*x),Nr,opt2);

            % check appropriateness equilibrium
            isequi=(norm(Nr.*(Ar+Br*Nr))<1e-6);
            isfeas=all(Nr>1e-6); % feasibility check
            jacr=diag(Ar+Br*Nr)+diag(Nr)*Br;
            isstab=(max(real(eig(jacr)))<0); % stability check
            chkcomm=isequi && isfeas && isstab;
        end
        comms(ic1,ic2)=struct('A',Ar,'B',Br,'N',Nr);
    end
end

%%% SECOND STEP: COMPUTE STABILITY MEASURES %%%

% for each community we compute the following set of stability measures:
% . asyresil: asymptotic resilience
% . varmean: mean-case variability, for 3 perturbation types alpha=0,1,2
%   (size 1x3)
% . varworst: worst-case variability, for 3 perturbation types alpha=0,1,2
%   (size 1x3)
% . varpert: variability measure for single-species perturbations, for 3
%   perturbation types alpha=0,1,2
%   (size nspx3, with nsp the number of species in the community)

stabs=struct('asyresil',cell(2,3), ...
             'varmean',cell(2,3), ...
             'varworst',cell(2,3), ...
             'varpert',cell(2,3));
	% structure to store stability values

for ic1=1:2,
    for ic2=1:3,
        A=comms(ic1,ic2).A;
        B=comms(ic1,ic2).B;
        N=comms(ic1,ic2).N;
        J=diag(A+B*N)+diag(N)*B;
        asyresil=-max(real(eig(J)));
        varmean=variability_meancase(N,J,[0 1 2]);
        varworst=variability_worstcase(N,J,[0 1 2]);
        nsp=length(N); % number of species in community
        varpert=zeros(nsp,3);
        for isp=1:nsp,
            E=zeros(nsp);
            E(isp,isp)=1;
            varpert(isp,:)=variability_perturbation(N,J,E,[0 1 2]);
        end
        stab=struct('asyresil',asyresil,'varmean',varmean, ...
            'varworst',varworst,'varpert',varpert);
        stabs(ic1,ic2)=stab;
    end
end
```
