## Supplementary material for "The inherent multidimensionality of temporal variability: How common and rare species shape stability patterns": Matlab code used to generate figures: variability_meancase.m

```
function Vmean=variability_meancase(N,A,al)
% function Vmean=variability_meancase(N,A,al) computes the mean-case
% variability metric given by equation (D7) in the paper.
% inputs:
% . N: vector of equilibrium abundances (size Sx1)
% . A: Jacobian matrix evaluated at equilibrium (size SxS)
% . al: perturbation type (scalar or vector)
% output:
% . Vmean: mean-case variability (same size as al)

Vmean=zeros(size(al));
nsp=length(N); % number of species
for ial=1:length(al),
    Vmean(ial)=trace(lyap(A,diag(N.^al(ial))))/nsp;
end

end
```
