## Supplementary material for "The inherent multidimensionality of temporal variability: How common and rare species shape stability patterns": Matlab code used to generate figures: variability_perturbation.m

```
function Vpert=variability_perturbation(N,A,E,al)
% function Vpert=variability_perturbation(N,A,E,al) computes the
% variability metric for a particular perturbation, given by equation
% (D5) in the paper.
% inputs:
% . N: vector of equilibrium abundances (size Sx1)
% . A: Jacobian matrix evaluated at equilibrium (size SxS)
% . E: perturbation covariance matrix (size SxS)
% . al: perturbation type (scalar)
% output:
% . Vpert: perturbation-specific variability (scalar)

Vpert=zeros(size(al));
for ial=1:length(al),
    D=diag(N.^(al(ial)/2));
    E=E/trace(E);
    Vpert(ial)=trace(lyap(A,D*E*D));
end

end
```
