## Supplementary material for "The inherent multidimensionality of temporal variability: How common and rare species shape stability patterns": Matlab code used to generate figures: variability_worstcase.m

```
function Vworst=variability_worstcase(N,A,al)
% function Vmean=variability_worstcase(N,A,al) computes the worst-case
% variability metric given by equation (D6) in the paper.
% inputs:
% . N: vector of equilibrium abundances (size Sx1)
% . A: Jacobian matrix evaluated at equilibrium (size SxS)
% . al: perturbation type (scalar or vector)
% output:
% . Vworst: worst-case variability (same size as scalar)

Vworst=zeros(size(al));
nsp=length(N); % number of species
for ial=1:length(al),
    Ahat=kron(A,eye(nsp))+kron(eye(nsp),A);
    Dhat=kron(diag(N.^(al(ial)/2)),diag(N.^(al(ial)/2)));
    Vworst(ial)=norm(inv(Ahat)*Dhat);
end

end
```
